# Local structural preference maps encode transferable protein-interface energetics

**DOI:** 10.64898/2026.09.18.752156

**Authors:** Maximilian Kienlein, Filipe Menezes, Phillip Grass, Stefan Munker, Till Siebenmorgen, Grzegorz M. Popowicz

**Affiliations:** Institute of Structural Biology, Molecular Targets and Therapeutics Center, Helmholtz Munich, Ingolstädter Landstraße 1, Neuherberg, 85764, Germany; Oncera GbR, Am Klopferspitz 19a, Martinsried, Germany.; Khumbu.AI GmbH, Gabelsbergerstraße 62, Munich, Germany.

**Keywords:** protein–protein interactions, structural machine learning, molecular recognition, binding affinity, interface specificity

## Abstract

Relating the drivers of binding affinity to structural features remains difficult because affinity emerges from many weak, context-dependent interactions, while experimental affinity and mutation data are limited. Here, we ask whether a machine-learning model can learn energetically relevant interaction preferences directly from native structures, without affinity or mutation labels. We decompose interfaces into local structural motifs and learn the compatibility of each motif with its surrounding molecular environment. Applied across a protein surface, these models yield target preference maps (TPMs), spatial fields of local interaction compatibility. Trained exclusively on native antibody-antigen structures without affinity or mutation labels, TPMs recover charged, aromatic and backbone-mediated recognition across peptide-protein and other non-antibody interfaces. On an out-of-domain SKEMPI benchmark, wild-type TPM scores discriminated mutation-sensitive positions (AUC 0.643, n=1323), while TPM-based substitution scores ranked alternative amino acids comparably to FoldX and Rosetta Flex ΔΔ*G*. These results show that native interface structures can supervise learning of transferable, energetically relevant molecular preferences without direct thermodynamic labels.

---

Protein–protein interactions underlie biological recognition and regulation, yet predicting how interface structure determines binding remains a major challenge [1–3]. Atomic structures reveal the geometric and chemical complementarity of bound complexes at high resolution, suggesting that molecular interaction preferences should, in principle, be inferable from the arrangement of atoms at an interface. Recent advances in structure prediction and generative protein design have made it increasingly possible to generate protein complexes with plausible interaction geometries [4–6]. Structural plausibility, however, does not establish whether an interface is energetically favourable, which positions are critical for binding, or which sequence changes will strengthen or weaken an interaction. Distinguishing among plausible interfaces and prioritizing beneficial mutations, therefore, remain central problems in rational interface optimization and protein design [7, 8]. A central difficulty is that protein binding is not determined by isolated contacts, but by the collective effect of many weak and context-dependent interactions distributed across an extended surface. These energetic contributions, however, are not distributed uniformly across the interface, as individual hot-spot residues can contribute disproportionately to binding [9].

Compared with small-molecule binding sites, protein interfaces are typically larger, more solvent-exposed and more conformationally heterogeneous [1, 10, 11]. The resulting binding free energy reflects a balance of hydrogen bonding, electrostatics, van der Waals packing, desolvation, water-mediated interactions and entropic contributions [12, 13]. The contribution of an individual chemical group therefore depends not only on its identity, but on its relative orientation, neighbouring residues and solvent environment. For example, a hydroxyl group may be strongly favoured in one local context and disfavoured only a few Å away.

These properties create complementary limitations for physics- and data-driven approaches. Physics-based methods can provide estimates of mutation effects with considerable atomic detail, but commonly require construction of every mutant structure, conformational sampling and energy relaxation. They remain sensitive to the input structure and force-field assumptions [10, 14]. Machine-learning approaches can instead learn structure-energy relationships directly from experimental measurements, but structures paired with affinity or mutation labels remain sparse and noisy relative to the enormous chemical and structural diversity of protein interfaces. Recent analysis of antibody-antigen ΔΔ*G* prediction showed that the apparent performance can deteriorate substantially under stringent generalization tests and suggested that orders of magnitude more diverse measurements may be required for robust prediction of mutation effects [15]. Experimental datasets are additionally shaped by assay dynamic range and experimental selection, contributing to bias in such training data. For example, weak or non-binding variants can be difficult to quantify precisely, and highly disruptive mutations may fall outside the measurable affinity range. Related proteins or structurally similar interfaces occurring across training and evaluation sets can further inflate apparent generalization [16, 17].

These limitations motivate an alternative question: can energetically relevant interaction preferences be learned from the much larger structural record of successful molecular recognition, without requiring affinity or mutation labels?

We reasoned that native protein interfaces contain such information at the level of recurring local molecular environments. Rather than learning a direct mapping from an entire complex to a binding free energy, we represent protein structures as collections of local interaction motifs, termed “shards”. This representation pools examples of equivalent chemistry across residues by defining motifs below the level of amino-acid identity, providing a less fragmented training signal (Fig. 1a). For each distinct shard type, separate models predict a compatibility score from the spatial distribution of target-atom arrangements commonly associated with that specific motif in experimentally observed interfaces. Importantly, the models are never presented with complete interfaces as training examples, since each prediction is conditioned only on the local environment surrounding the respective interaction motif. This local-only formulation leaves little opportunity to memorize complete interface identities and provides an explicit inductive bias towards generalization of recurring molecular interaction patterns that can, in principle, be reused across structurally unrelated complexes.

**Fig. 1.**
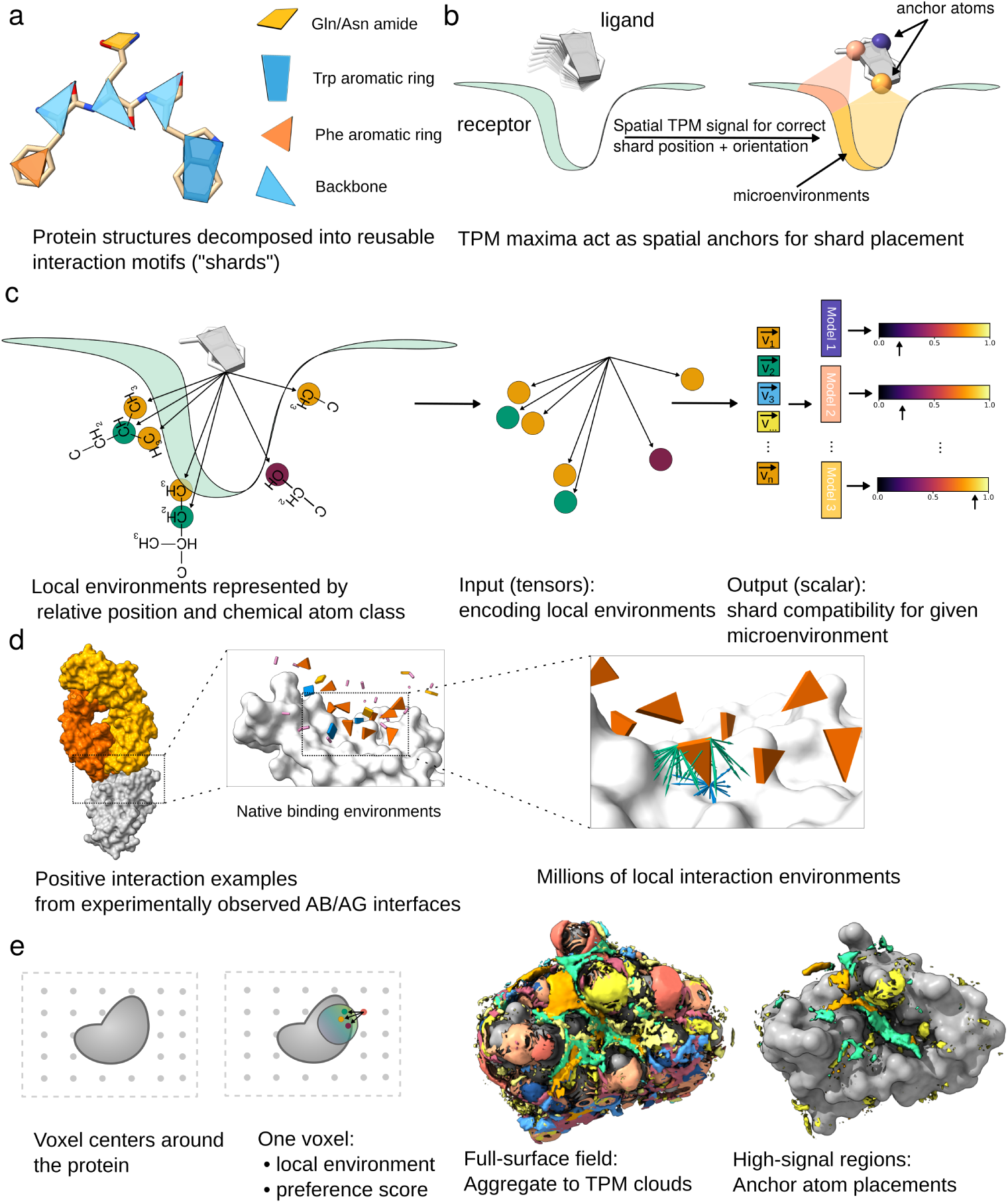
Target preference maps represent protein interfaces as spatial fields of local interaction compatibility. **a**, Protein structures are decomposed into rigid, reusable local interaction motifs, termed shards. These are defined below the level of amino-acid identity and encompass both side-chain and backbone groups. **b,** Trained shard-specific models are used to infer the compatibility of candidate anchor placements around the target surface, with high-scoring positions indicating locally compatible environments. Constituent anchor atoms jointly define the position and orientation of the complete motif (Extended Data Table 2 for full shard and anchor definitions). **c,** For each native shard placement, neighbouring target atoms define a local structural environment encoded by atom-class labels and position vectors relative to the anchor atom. **d,** Native antibody–antigen environments provide positive training examples. The decomposition of interfaces into microenvironments generates a multitude of training examples per interface. **e,** Evaluating the trained models across three-dimensional space yields target preference maps (TPMs), continuous fields identifying preferred positions for specific interaction motifs around the target surface.

Aggregating these local compatibility predictions in three-dimensional space yields target preference maps (TPMs), extending a representation previously introduced for small-molecule recognition [18]. TPMs are continuous spatial fields describing where defined chemical groups are supported by the local structural environment of a protein surface. Electrostatics, packing, solvation and other contributions are not prescribed as independent energetic terms. Rather, their combined consequences are inferred from the statistics of the molecular contexts in which particular interaction motifs are observed. A protein interface can, therefore, be interrogated as an assembly of independently learned local preferences rather than evaluated only as a single global object.

We selected antibody-antigen complexes as the training space because they provide a large and structurally diverse collection of interfaces shaped by highly specific molecular-surface recognition [19]. TPM training uses only native complex structures. The model receives no experimental binding affinities, mutation measurements, or ΔΔ*G* labels as input. Because prediction is local rather than global, transfer does not require preservation of the global topology of the training complexes. To ensure that measured transfer reflects genuine generalization rather than similarity between training and evaluation structures, we keep training and evaluation strictly separate and additionally test generalization to interface classes outside the training domain.

Here, we show that TPMs recover specific interaction preferences across structurally diverse protein interfaces, including charged, aromatic and backbone-mediated recognition. When summed across complete interfaces, TPM scores show a modest association with absolute binding affinity, indicating that local structural compatibility captures part, but not all, of the determinants of binding free energy. More directly, TPMs capture two quantities shaped by local interface context: the sensitivity of an interface position to sequence perturbation and the relative compatibility of alternative substitutions at that position. Native TPM scores identify positions at which mutations strongly perturb experimental binding in a single rapid scan of the wild-type structure, while TPM-derived substitution scores recover the relative ordering of mutations within individual pockets. On the independent SKEMPI benchmark, this ranking performance is comparable to FoldX and Rosetta Flex ΔΔ*G*, methods developed specifically for mutation-effect prediction [20, 21]. This is despite TPMs not using either backbone remodeling or energetic relaxation of mutant complexes, both of which dominate the computational cost of these methods. Together, these results establish locally learned structural preference fields as an interpretable and transferable representation of protein-interface energetics, and suggest that such fields could provide explicit local guidance for interface optimization and generative protein design.

## 1 Results

### 1.1 A structural prediction framework for protein interface specificity

We developed a target preference map framework to learn local structural determinants of protein-interface specificity from experimentally resolved complexes. Rather than representing a protein-protein interaction as a single global binding event, the framework decomposes interfaces into local interaction motifs. Each motif comprises a shard on the binder side and its environment, defined by the spatial arrangement of receptor atoms surrounding it (Fig. 1a, c). Shards are rigid chemical groups obtained by decomposing residues into fragments with fixed internal geometry, defined below the level of amino-acid identity and encompassing both side-chain and backbone interaction motifs (Extended Data Table 2). This decomposition exploits the small and chemically constrained residue alphabet of proteins.

Each shard contains a defined set of anchor atoms whose fixed internal geometry specifies its position and orientation (Fig. 1b). Separate preference fields are learned for these anchors.

For each native shard placement, neighbouring atoms on the opposing receptor define the corresponding target environment. These neighbours are encoded by their chemical atom-class labels and relative position vectors with respect to the anchor atom, retaining both the chemical composition and local geometry of the interaction environment (Fig. 1c, Methods 3.3). During training, native antibody–antigen interfaces provide positive examples of compatible molecular environments (Fig. 1d). Negative examples combine environments sampled from non-interface surface regions with native environments corresponding to other anchors, including the remaining anchors of the same shard (Extended Data Fig. A1). Because each classifier must distinguish its anchor from the others of the same shard, the individual anchor fields become spatially localized. This allows orientation to be encoded through their spatial relationship rather than predicted as an independent variable. The position and orientation of the complete motif are recovered from their joint preferences.

At inference, the trained models are used to score candidate anchor placements around a target protein, assigning each placement a local compatibility score (Fig. 1b). Aggregating these shard-level predictions in three-dimensional space yields target preference maps (TPMs), continuous fields indicating where specific interaction motifs are compatible with the local target environment (Fig. 1e). TPMs therefore provide an interpretable intermediate representation of molecular recognition. Rather than returning only a binary interaction prediction or a single global interface score, they expose the spatial distribution of residue- and atom-specific preferences around a protein surface. The learned preferences can therefore be tested directly against the functional groups observed in experimentally resolved complexes and examined for whether the same local interaction signals also encode the energetic effects of interface mutations.

In the following sections, we first test whether interaction preferences learned by models trained exclusively on antibody–antigen structures generalize to held-out antibody-antigen complexes and to structurally distinct peptide-protein and non-antibody protein-protein interfaces. We then examine whether these preferences carry local energetic information and, despite the absence of affinity or mutation labels during training, transfer to experimentally measured mutation effects.

### 1.2 Interface predictions generalize across structurally diverse complexes

In order to test whether local preferences transfer, it is necessary to ensure that measured performance reflect genuine generalization rather than similarity to the training structures. We therefore separated each evaluation set from training according to its benchmark design, controlling antibody-antigen redundancy at the CDR-sequence level and using the peptide-protein and SKEMPI benchmarks as external tests outside the antibody-antigen interaction class (Table 1).

**Table 1.** Dataset provenance across model training and evaluation. Antibody-antigen redundancy was controlled by clustering complementarity-determining-region (CDR) sequences at 70% identity, with complete clusters assigned to train, validation or test splits. Peptide-protein and SKEMPI benchmarks provide external tests outside the antibody-antigen interaction class used for TPM training. For SKEMPI, antibody-antigen and TCR/pMHC complexes were excluded.

| Dataset / benchmark | Role | Struct. | Evaluation scale | Main use |
| --- | --- | --- | --- | --- |
| SAbDab antibody-antigen | Training | 5,110 | 1,194,403 shard env.<sup>a</sup> | TPM model training |
| SAbDab antibody-antigen | Validation | 498 | 111,799 shard env. | Model selection / thresholds |
| SAbDab antibody-antigen | Held-out test | 683 | 164,476 shard env. | In-distribution evaluation |
| Independent peptide-protein | External test | 185 | 16,151 shard env. | Cross-class structural generalization |
| SKEMPI mutation benchmark | External test | 217 | 1,323 positions / 2,833 substitutions | Hotspot detection and substitution ranking |

<sup>a</sup> Positive shard environments retained after the microenvironment validity filter (36 anchor classes, Methods 3.3). Negatives are resampled at every training epoch and are therefore not included in these counts.

**Table 2.** Anchor classes used throughout this study. Each anchor class is defined by one or more residue–atom combinations that provide equivalent anchor positions for a single classifier, and multiple anchor classes may jointly represent one chemical group (for example Arg-CZ and Arg-NH for the guanidinium). The labels correspond to the terminology used in the main figures and text. An asterisk indicates a residue-independent class where the specified atom is present.

| Shard anchors | Residue(s) | Anchor atom(s) |
| --- | --- | --- |
| BB-$C\beta$ | * | CB |
| BB-O | * | O |
| BB-H | * | H |
| Lys-NZ | Lys | NZ |
| Lys-CE | Lys | CE |
| Arg-CZ | Arg | CZ |
| Arg-NH | Arg | NH1, NH2 |
| Ser/Thr-O | Ser, Thr | OG, OG1 |
| Ser/Thr-$C\beta$ | Ser, Thr | CB |
| Cys-SG | Cys | SG |
| Cys-$C\beta$ | Cys | CB |
| Met-SD | Met | SD |
| Met-CE | Met | CE |
| Asp/Glu-C | Asp, Glu | CG, CD |
| Asp/Glu-O | Asp, Glu | OD1/OD2, OE1/OE2 |
| Asn/Gln-C | Asn, Gln | CG, CD |
| Asn/Gln-O | Asn, Gln | OD1, OE1 |
| Asn/Gln-N | Asn, Gln | ND2, NE2 |
| Phe-CG | Phe | CG |
| Phe-CE | Phe | CE1, CE2 |
| Tyr-OH | Tyr | OH |
| Tyr-CD | Tyr | CD1, CD2 |
| Trp-CG | Trp | CG |
| Trp-NE1 | Trp | NE1 |
| Trp-CZ3 | Trp | CZ3 |
| His-CG | His | CG |
| His-ND1 | His | ND1 |
| His-NE2 | His | NE2 |
| Pro-CA | Pro | CA |
| Pro-CG | Pro | CG |
| Pro-N | Pro | N |
| Gly-CA | Gly | CA |
| Ala-$C\beta$ | Ala | CB |
| Leu-CG | Leu | CG |
| Val-$C\beta$ | Val | CB |
| Ile-$C\beta$ | Ile | CB |

**Table 3.** Target-environment atom classes. Each neighbouring target atom is assigned to a chemical atom classes based on its atom name and parent residue, defining the atom-class of the local environment representation (Methods 3.6). These classes describe the chemical character of environment atoms and are distinct from the shard anchor classes of Extended Data Table 2.

| Chemistry | Atom(s) | Residue(s) |
| --- | --- | --- |
| Aliphatic carbon | CA, CB, CG, CD, CE | CA, CB in all residues; CG, CD, CE in ILE, LEU, VAL, PRO, THR, MET, LYS, ARG, GLN, GLU |
| Aromatic carbon | CG, CD, CE, CZ | PHE, TYR, TRP, HIS |
| Backbone amide nitrogen | N | all |
| Backbone carbonyl carbon | C | all |
| Hydroxyl oxygen | OG | SER, THR |
| Carboxylate oxygen | OD, OE | ASP, GLU |
| Aromatic hydroxyl oxygen | OH | TYR |
| Amide carbonyl oxygen | OD, OE | ASN, GLN |
| Amide nitrogen | ND, NE | ASN, GLN |
| Amide carbon | CG, CD | ASN, GLN |
| Carboxyl carbon | CG, CD | ASP, GLU |
| Indole nitrogen | NE | TRP |
| Thioether sulfur | SD | MET |
| Thiol sulfur | SG | CYS |
| Guanidinium carbon | CZ | ARG |
| Imidazole nitrogen | ND, NE | HIS |
| Guanidinium NE | NE | ARG |
| Charged amine nitrogen | NH, NZ | ARG, LYS |
| Backbone carbonyl oxygen | O | all |

Antibody-antigen interfaces provide a rich source of structurally resolved molecular recognition, but represent only one class of protein-protein interaction. Therefore, we asked whether local interaction preferences learned exclusively from antibody-antigen structures transfer to interfaces with different architectures and biological functions. To test this quantitatively, we compared shard-level discrimination on the held-out antibody-antigen test set with an independent peptide-protein test set held out from training and model selection (Fig. 2a). Across most anchor classes, discrimination transferred directly to the peptide-protein set, with peptide AUCs spanning approximately 0.55-0.90 and broadly tracking the corresponding held-out antibody-antigen values (Fig. 2a).

**Fig. 2.**
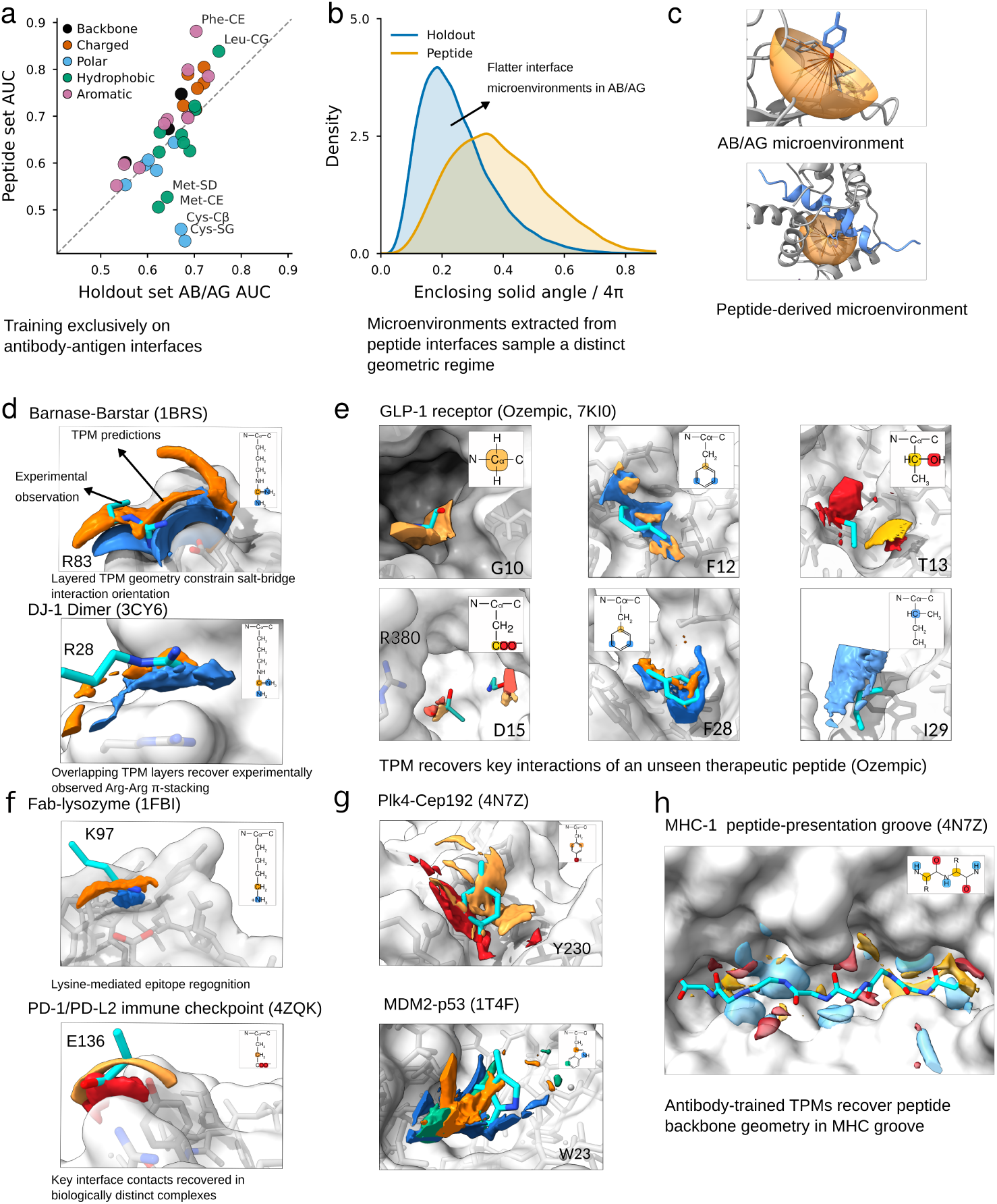
Interface predictions generalize across structurally diverse complexes. **a**, Shard-level AUCs on held-out antibody–antigen complexes and an independent peptide–protein test set. Models were trained exclusively on antibody-antigen structures. Evaluation used a stringent negative set composed of alternative native shard environments from the same test structures, excluding random non-interface surface environments (Methods 3.9). **b,** Enclosing solid-angle distributions show that peptide-derived microenvironments occupy a more spatially enclosed geometric regime than antibody–antigen environments. **c,** Representative local environments illustrating the enclosing solid-angle measure used in **b**. For **d–h**, TPMs were calculated using only the receptor structure. The corresponding ligand-side functional groups (in cyan) shown from the solved complexes were not used to generate the predictions. **d,** TPMs resolve distinct interaction geometries of the same functional group. In barnase–barstar (PDB 1BRS), the nitrogen-centred arginine field lies adjacent to barstar Asp39 while the C*ζ* field is displaced behind it, reconstructing the native salt-bridge orientation of barnase Arg83. In the DJ-1 homodimer (PDB 3CY6), the nitrogen- and C*ζ*-centred fields are instead nearly coplanar with the opposing guanidinium group, reproducing the stacked Arg28–Arg48 geometry. The anchor-field geometry therefore reveals how the same arginine motif can support fundamentally different interactions, from conventional charge-complementary salt bridging to like-charged guanidinium *π*-stacking. **e,** TPMs at important semaglutide positions implicated by mutagenesis [24]. **f,** Preference fields recover lysozyme Lys97 in the Fab-lysozyme interface (PDB 1FBI) and PD-1 Glu136 in the PD-1/PD-L1 immune-checkpoint complex (PDB 4ZQK). **g,** Aromatic TPMs recover Cep192 Tyr230 and p53 Trp23 positions. The less confined Trp23 prediction is consistent with pre-dominantly hydrophobic recognition imposing weaker spatial constraints compared to e.g. a salt bridge. **h,** Backbone TPMs trace the ovalbumin peptide through an MHC-I binding groove, showing transfer to extended backbone-mediated recognition in addition to side-chain-specific contacts.

This evaluation uses a deliberately stringent negative set. For each shard model, positives are held-out native environments of that shard. The negatives are taken from the other native environments of the same test structures, including chemically related but distinct shard classes and alternative anchors of the same functional group. Random non-interface surface environments, although used during training, are excluded from evaluation. Consequently, an AUC of 0.5 corresponds to chance discrimination among alternative native molecular environments rather than between an interface environment and bulk solvent.

This transfer occurred despite a measurable geometric shift between the antibody–antigen and peptide-protein regimes. Local environments extracted from peptide–protein complexes occupied a more spatially enclosed regime than antibody–antigen environments, as quantified by their enclosing solid angles (Fig. 2b,c). This is consistent with the more groove- and pocket-like recognition geometry frequently encountered in peptide-protein complexes. The peptide test set, therefore, probes the model in local environments that differ systematically from the antibody-antigen training distribution. Together with the shard-level discrimination in Fig. 2a, this shows that the learned preferences transfer not only across protein families but also to a distinct local structural regime.

Performance varied across motif classes. Aromatic and backbone-centred motifs generally retained strong discrimination, whereas cysteine- and methionine-associated shards were among the clearest low-performing classes. Reduced cysteine performance is consistent with the limited representation of cysteine interaction environments in the antibody-antigen training data. In particular, interfacial disulfide geometries are essentially absent, leaving this recognition mode unsupported. The origin of the reduced methionine performance remains unclear.

We next asked whether this quantitative transfer corresponds to chemically meaningful spatial predictions in individual complexes outside the antibody-antigen training regime. Whereas the shard-level AUC analysis evaluates discrimination across complete test sets, the following examples directly compare predicted TPM fields with experimentally observed functional-group positions in solved structures (Fig. 2d–h). For this we calculated the TPM field using only the receptor structure. The corresponding ligand-side functional group, shown from the solved complex, was not used to generate the prediction and is only included as a reference for evaluating the predictions. The TPM clouds represent regions of local compatibility at a selected contour threshold, with higher contour levels progressively isolating the strongest predicted anchor positions.

Arginine provides a particularly clear example of how these fields encode interaction geometry rather than simply chemical identity. In the barnase–barstar complex (PDB 1BRS), the nitrogen-centred preference field lies adjacent to barstar Asp39, while the C*ζ* field is displaced by approximately one guanidinium bond length, reconstructing the native salt-bridge orientation of barnase Arg83. By contrast, in the DJ-1 homodimer (PDB 3CY6), the nitrogen- and C*ζ*-centred fields are nearly coplanar with the opposing guanidinium group, reproducing the stacked Arg28–Arg48 geometry (Fig. 2d). Such close-range guanidinium *π*-stacking is a recurring structural motif in proteins, with direct parallel Arg-Arg contacts observed for approximately 3.6% of arginine side chains despite the electrostatic repulsion expected between like-charged groups [22]. Its recovery by the arginine TPM illustrates how recurring, non-obvious interaction patterns present in native protein structures can emerge directly from structural training data. Thus, rather than identifying only regions favourable to positive charge, the relative arrangement of the TPM fields reveals the distinct interaction geometries of the same functional group.

Residue-specific TPMs also recovered interaction geometries across diverse peptide and protein interfaces. In the semaglutide-GLP-1 receptor complex (PDB 7KI0) [23], we examined ligand positions previously identified by mutagenesis as important for GLP-1 receptor binding [24]. TPMs recovered several of these experimentally resolved interaction geometries, including a strongly activity-sensitive Asp15-Arg380 salt bridge (Fig. 2e). Agreement was likewise observed for lysozyme Lys97 in the Fab-lysozyme complex (PDB 1FBI) and PD-1 Glu136 in the PD-1/PD-L1 immune-checkpoint complex (PDB 4ZQK, Fig. 2f), and for Cep192 Tyr230 in the Plk4-Cep192 complex (PDB 4N7Z) and Trp23 of an optimized p53-derived peptide bound to MDM2 (PDB 1T4F, Fig. 2g). Importantly, transfer was not limited to side-chain recognition. Backbone TPMs traced the experimentally observed path of the ovalbumin peptide through the MHC-I H-2Kb binding groove (PDB 1VAC), recovering extended backbone-mediated recognition across the peptide interface (Fig. 2h).

Together, the quantitative and structural analyses show that locally learned antibody-antigen interaction preferences transfer across substantial changes in interface architecture, protein family and chemical context. This is consistent with the local representation capturing reusable spatial constraints on molecular recognition rather than antibody-specific interface architecture.

### 1.3 TPMs resolve residue-level interaction grammar and identify mutation-sensitive interface positions

We next asked what level of molecular specificity these fields encode. A local environment need not map to a single shard identity, because related groups may satisfy the same structural constraints. This degeneracy was directly visible in individual interfaces (Fig. 3a,b). In barnase-barstar, Phe- and Trp-compatible fields overlap in the hydrophobic pocket at position 38 (Fig. 3a). At SARS-CoV-2 RBD position 501, the wild-type Asn environment also supports a Tyr-compatible field that overlaps the experimentally observed N501Y side chain after alignment to an independently solved mutant complex (Fig. 3b). In these cases, TPM overlap represent alternative structurally compatible solutions rather than model confusion.

**Fig. 3.**
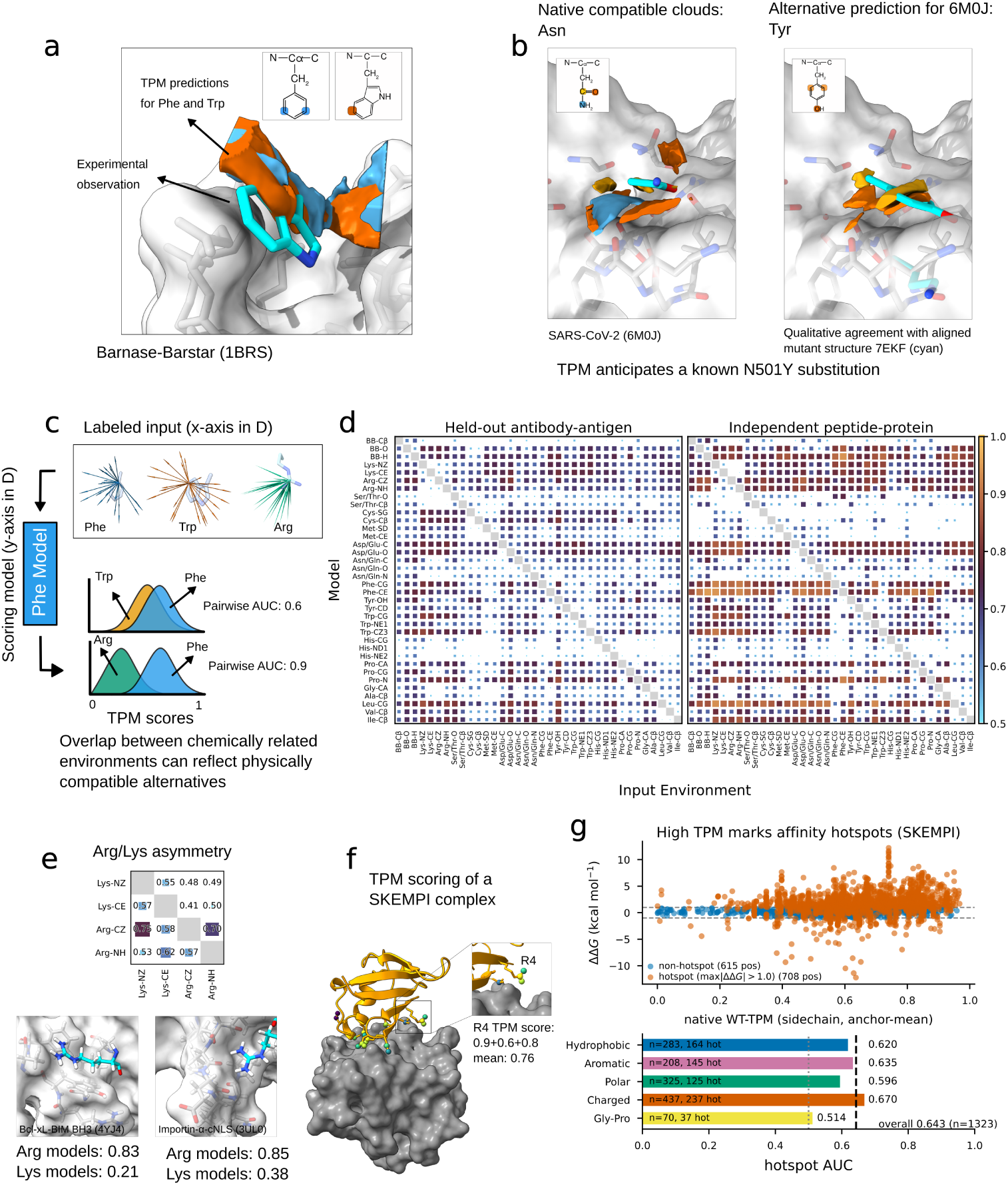
TPMs resolve residue-level interaction grammar and identify mutation-sensitive interface positions. **a, b**, Overlapping preference fields can represent alternative structurally compatible residue identities. **a,** Pheand Trp-compatible TPM fields overlap in a hydrophobic pocket of the barnase-barstar interface, illustrating aromatic degeneracy. **b,** At SARS-CoV-2 RBD position 501, the wild-type receptor environment supports both the native Asn-compatible field and a Tyr-compatible field; after alignment to an independently solved N501Y complex (PDB:7EKF), the latter overlaps the experimentally observed mutant tyrosine. Together, these examples illustrate that a single local environment can support multiple structurally compatible residue identities. **c,** Schematic of pairwise shard anchor discrimination. For each row anchor, score distributions are compared between native environments of the same anchor and native environments belonging to a second shard anchor class. **d,** Pairwise AUC matrices on held-out antibody–antigen and independent peptide–protein interfaces. High AUC indicates distinct environmental preferences, whereas reduced off-diagonal AUC indicates overlapping compatibility and need not correspond to model confusion. **e,** Asymmetric discrimination between lysine and arginine environments. Arginine models distinguish their native environments from lysine environments more strongly than lysine models do in reverse (mean pairwise AUC 0.62 versus 0.47), consistent with the stronger geometric constraints imposed by the planar guanidinium group. Structural examples illustrate guanidinium-specific recognition through planar packing and directional hydrogen bonding. **f,** Native TPM scoring of a representative SKEMPI interface. Wild-type interface positions are scored directly from the native complex without introducing the experimentally measured mutations. **g,** Native TPM scores identify mutation-sensitive interface positions. Across 1323 SKEMPI interface positions, native scores discriminate mutation-critical positions, defined by at least one measured mutation with |ΔΔ*G| >* 1 kcal,mol*^−^*^1^, from tolerant positions (overall AUC 0.643). Discrimination is retained across hydrophobic, aromatic, polar and charged native residue categories (AUCs 0.620, 0.635, 0.596 and 0.670, respectively).

To quantify this systematically, we computed pairwise AUC matrices between anchor classes. Each shard model was evaluated on native environments belonging either to its own class or to a second class (Fig. 3c). High pairwise AUC indicates the model separates the two distributions, whereas values near 0.5 indicate overlapping compatibility. Pairwise matrices obtained independently on held-out antibody-antigen complexes and on the PDBbind-derived peptide-protein test set showed similar off-diagonal structure (Fig. 3d), indicating that these residue-level relationships persist across interaction classes.

Chemically related residues were nevertheless not collapsed into single coarse categories. Lysine and arginine, for example, carry the same formal charge but displayed an asymmetric pairwise relationship. The arginine models separated their native environments from lysine environments more strongly than the lysine models separated theirs from arginine (mean pairwise AUC 0.62 versus 0.47, with several lysine-versus-arginine comparisons falling below 0.5; Fig. 3e). This asymmetry has a structural interpretation. The flexible lysine terminal amine presents a relatively compact charge and hydrogen-bond donor group with weak orientational constraints. The arginine guanidinium group has greater reach and presents a larger charge-bearing and hydrogen-bond donor surface, while its planar geometry supports more specific directional hydrogen-bonding and stacking arrangements. Many arginine environments are therefore scored as acceptable by the less restrictive lysine models, whereas the arginine models impose more specific geometric requirements. Structural examples illustrate these modes through planar packing and multi-hydrogen-bond arrangements with backbone acceptors (Fig. 3e). TPMs therefore distinguish interaction motifs not only by broad physicochemical properties such as charge or aromaticity, but also by the geometric constraints imposed by individual functional groups.

To test whether these local compatibility signals correspond to interaction energetics, we turned to experimental mutation data. We evaluated TPMs on SKEMPI (Table 1), a demanding out-of-domain test of measured binding free-energy changes (ΔΔ*G*) on non-antibody complexes, assembled independently of the training data. We first asked whether the unperturbed interface itself reveals which positions are most sensitive to sequence change. Each mutated interface position was scored directly from its wild-type structure, without introducing any of the measured substitutions (Fig. 3f). Positions were classified as mutation-critical when at least one measured substitution produced |ΔΔ*G| >* 1 kcal mol*^−^*^1^. Across 1323 interface positions, native TPM scores discriminated mutation-critical from tolerant sites with an AUC of 0.643 (Fig. 3g). Discrimination was retained within hydrophobic, aromatic, polar and charged residue classes, with AUCs of 0.620, 0.635, 0.596 and 0.670, respectively. Because these scores are calculated from the unmutated interface alone, the native local environment already contains information about how strongly a position responds to subsequent sequence perturbation.

As an independent mechanistic check on whether this structural signal reflects interaction energetics, we compared TPM score differences for matched phenylalanine/tyrosine substitutions in the peptide-protein set with quantum-mechanical interaction-energy differences (Extended Data Fig. A3, Methods 3.12). A modest association appeared only when the QM calculation included implicit solvation (*ρ* = *−*0.28, *n* = 182), not for gas-phase energies (*ρ* = +0.12). The mutation-sensitivity result on experimental ΔΔ*G* and the solvation dependence of this control converge, indicating that the native interface structure encodes energetic contributions beyond the explicit pairwise terms of a physics-based force field, most directly the desolvation cost carried by the native, solvated structures TPMs learn from.

As an even coarser test, we asked how much energetic signal survives the simplest possible aggregation, an unweighted sum of TPM compatibility scores across an entire interface. Even this global sum showed a modest but consistent positive association with absolute binding affinity on the independent peptide-protein set (Pearson *r* = 0.33, Spearman *ρ* = 0.30, *n* = 173, Extended Data Fig. A2), which spans a wide affinity range and provides a more stringent test than the smaller SARS-CoV-2 RBD set, where only a weaker, non-significant trend in the same direction was observed. Notably, a physics-based MM/GBSA baseline computed on the same peptide-protein complexes shows no stronger association with experimental affinity (Extended Data Fig. A2). That an affinity-independent representation retains measurable information about absolute binding strength, without any weighting or calibration, indicates that recurring local structural preferences capture part of the energetic determinants of binding.

Together, these analyses show that TPMs encode a residue-level interaction grammar in which related motifs can occupy overlapping preference space while retaining functional-group-specific geometric constraints, and that this same local representation carries energetic information, from single-position mutation sensitivity up to a modest whole-interface affinity signal, without ever being trained on affinity or mutation labels.

### 1.4 TPM scores rank interface substitutions without relying on affinity data

Having identified mutation-sensitive interface positions, we next asked whether TPMs could rank alternative substitutions within individual pockets. Using the same out-of-domain SKEMPI benchmark introduced above, we restricted the analysis to positions with at least 15 measured substitutions (*n* = 33, Methods 3.13) and compared the predicted ordering of alternative amino-acid identities with the experimentally measured ΔΔ*G* series.

For each pocket, TPM fields for all candidate substitutions were generated from the native receptor environment and candidate side-chain conformations were placed by rotamer and pose search (Fig. 4a). Each pose received a raw fit score quantifying its agreement with the corresponding anchor fields, with higher values indicating better fit. The highest-scoring pose defined the raw fit score for that substitution. For comparison with experimental, FoldX and Flex ΔΔ*G*, this score was sign-inverted so that lower values denote more favourable substitutions across all three methods.

**Fig. 4.**
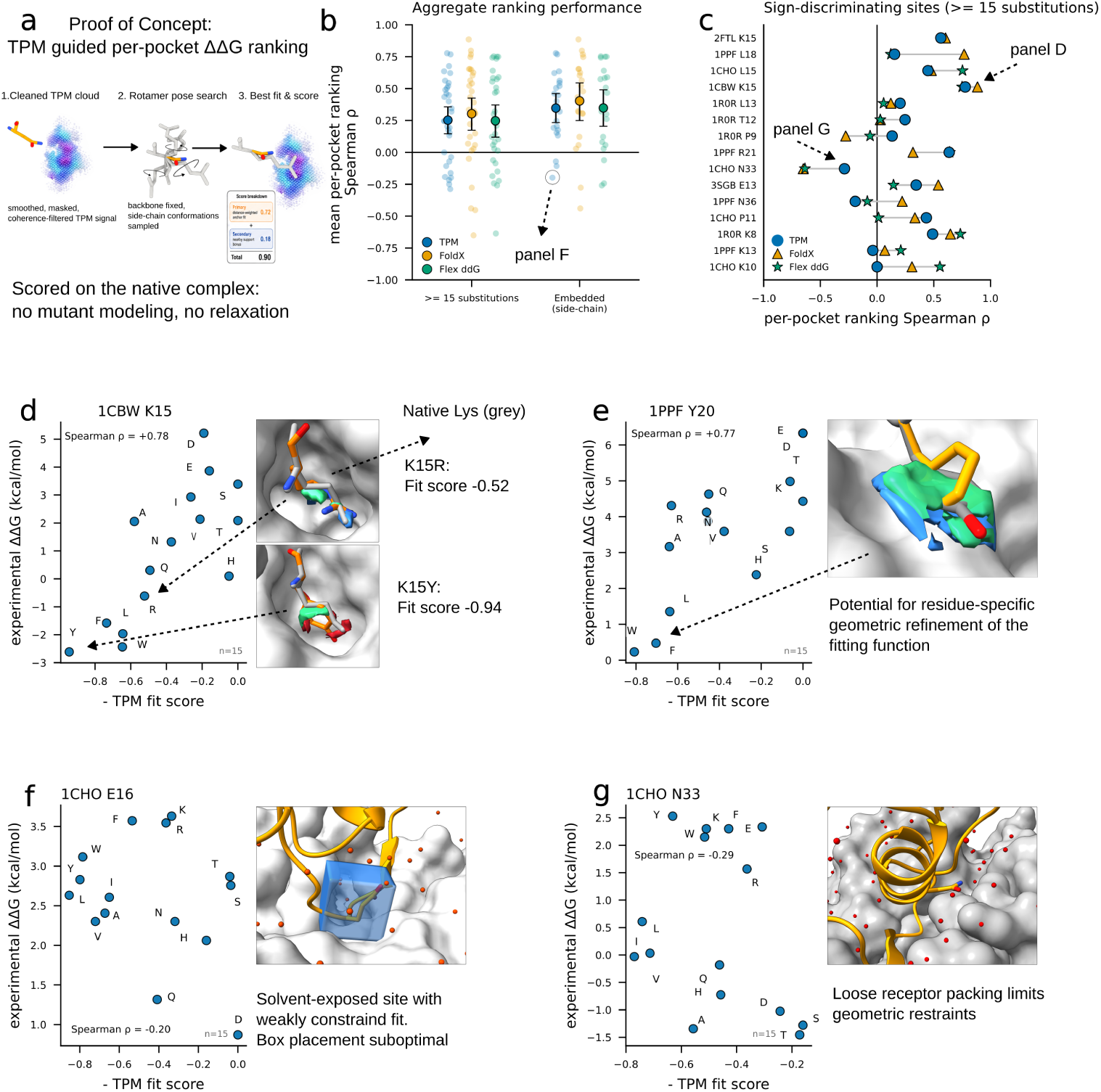
TPM ranks interface substitutions by their effect on binding without affinity training. **a**, Per-pocket substitution-ranking workflow. TPM fields are generated from the native receptor environment, and candidate side-chain conformations are placed by rotamer and pose search. Each pose receives a raw fit score according to its agreement with the corresponding anchor fields, with higher values indicating better fit The highest-scoring pose defines the raw fit score for each substitution. For quantitative comparison with experimental, FoldX and Flex ΔΔ*G*, this value is sign-inverted to obtain the TPM substitution score, such that lower values denote more favourable substitutions across all three methods. **b,** Aggregate per-pocket ranking performance (mean Spearman *ρ* between predicted and experimental ΔΔ*G* within each pocket) for TPM compared with FoldX and Flex ΔΔ*G* on the same pockets, for pockets with *≥* 15 substitutions (*n* = 33) and for the embedded-pocket subset (side-chain embeddedness *≥* 0.04; *n* = 21). Points are individual pockets. **c,** Per-pocket Spearman *ρ* across all sign-discriminating pockets (*≥* 15 substitutions, *n* = 15), TPM versus FoldX and Flex ΔΔ*G*; unlike **b**, this panel is not restricted to the embedded subset. **d–g,** Representative pockets showing the sign-inverted TPM substitution scores for individual amino-acid identities. Lower values denote more favourable predicted substitutions. **d,** 1CBW K15 and **e,** 1PPF Y20: geometrically well-defined sites where TPM ranks substitutions in agreement with experiment (*ρ* = +0.78 and +0.77). In **e**, the Phe TPM field overlaps the native Tyr planar ring, whereas the fixed C*α*/C*β* pose search places the fitted ring at a slight tilt. **f,** 1CHO E16 and **g,** 1CHO N33: solvent-exposed or loosely packed sites where weak geometric constraint limits ranking accuracy (*ρ* = *−*0.20 and *−*0.29).

Across qualifying SKEMPI pockets, TPM recovered the experimental mutation ordering with a mean per-pocket Spearman *ρ* of 0.25, compared with 0.30 for FoldX and 0.25 for Flex ΔΔ*G* on the same pockets (Fig. 4b,c). TPM therefore achieved similar mean ranking performance to two methods developed specifically for mutation-effect prediction, while using preference fields learned from native structures and evaluating substitutions without backbone remodeling or energetic relaxation of mutant complexes.

Ranking performance depended strongly on the geometric constraint of the mutated site. For TPM, mean Spearman *ρ* increased from 0.25 across all qualifying pockets to 0.35 for the 21 embedded pockets, defined by a side-chain embeddedness of *≥* 0.04, where side-chain embeddedness is the occluded solid-angle fraction of the native side-chain anchor atoms (Fig. 4b, Extended Data Fig. A4, Methods 3.13). The same trend was observed for FoldX (0.30 *→* 0.40) and Flex ΔΔ*G* (0.25 *→* 0.35), suggesting that local geometric embedding is a common axis of mutation-ranking predictability rather than a TPM-specific effect. Pockets in which the scored position directly contacted a partner histidine tended to show poorer TPM ranking performance (Extended Data Fig. A4), consistent with uncertainty in histidine protonation and tautomer assignment inherited from the structural training data (Methods 3.2).

Per-pocket correlations showed substantial site-to-site heterogeneity for all three methods (Fig. 4c), and individual pockets illustrate its structural basis (Fig. 4d–g). At 1CBW K15 and 1PPF Y20, where the substituted side chain is strongly constrained by the surrounding interface, TPM ranked chemically diverse substitutions in close agreement with experiment (*ρ* = +0.78 and +0.77, respectively). By contrast, agreement degraded at solvent-exposed or loosely packed sites, yielding *ρ* = *−*0.20 at 1CHO E16 and *ρ* = *−*0.29 at 1CHO N33. Per-pocket zoom-in views for a representative subset are shown in Extended Data Figs. A5–A25. These cases suggest that the current pose-based readout is most informative where the immediate structural environment strongly constrains alternative side-chain placements.

The spatial fields additionally reveal where errors arise in converting these preferences into a scalar substitution score. At 1PPF Y20, for example, the phenylalanine preference field recovers the native aromatic ring in its experimentally observed planar orientation (Fig. 4e), yet the rotamer and pose search fixes the C*α*/C*β* frame and places the candidate ring at a slight tilt relative to this preferred field. Thus, even where the learned field captures the native interaction geometry, translating it into a single substitution score introduces an additional approximation. The present pose-based fit should therefore be viewed as a simple readout of the underlying preference fields rather than an optimized mutation-energy estimator.

This transfer to an out-of-domain mutation benchmark, at performance comparable to dedicated ΔΔ*G* methods on the same pockets, supports the interpretation that the learned preference fields contain transferable information relevant to protein-interface energetics.

## 2 Discussion

This study asks whether experimentally resolved native interfaces can themselves supervise learning of molecular interaction energetics without affinity or mutation labels. It finds that they largely can, reaching mutation-ranking performance comparable to established ΔΔ*G* methods at substantially lower per-mutation computational cost, since no mutant structures are constructed or relaxed. Target preference maps realize this idea by decomposing resolved interfaces into local interaction motifs and learning the structural environments in which each motif is compatible. Rather than collapsing an interface to a single score, TPMs expose where defined interaction motifs are favoured around a protein surface, making the learned preferences directly interpretable in terms of atomic interaction geometry.

A central finding is that interaction preferences learned from antibody-antigen complexes transfer beyond the training domain. TPMs retained shard-level discrimination on a peptide-protein test set, recovered side-chain and backbone placement in peptide-binding grooves, and reproduced local interaction geometry in non-antibody protein-protein complexes. Antibody-antigen structures therefore contain reusable examples of molecular recognition, despite their particular evolutionary and structural context, that support learning of interaction preferences beyond antibodies. This transferability is consistent with recent evidence that antibodies can recapitulate interaction patterns used by cognate non-antibody ligands, suggesting that target surfaces can constrain molecular recognition toward convergent interaction solutions [25]. These examples also illustrate the interpretability of the representation, since the relative arrangement of anchor fields resolves orientation-specific interaction modes, including both conventional charge-complementary interactions and less intuitive geometries such as guanidinium stacking. The pairwise residue analysis further shows that the learned preferences are not limited to broad physicochemical categories. TPMs capture both degeneracy, where chemically related residues satisfy similar local constraints, and discrimination, where residues of similar charge or hydrophobicity remain distinguishable through functional-group-specific geometry.

The clearest evidence that these structural preferences carry binding-relevant information comes from their transfer to experimental energetic data. Native TPM scores identified mutation-sensitive interface positions in the out-of-domain benchmark, and TPM-derived substitution scores recovered the relative ordering of alternative amino acids within individual pockets, with mean per-pocket performance comparable to FoldX and Rosetta Flex ΔΔ*G* on the same substitutions. This is notable because TPMs were trained solely on native structures and the substitution readout requires neither construction nor relaxation of mutant complexes. Even an unweighted global sum of TPM scores retained a noticeable association with absolute binding affinity across the independent peptide-protein set, indicating that some information about overall binding strength survives the crudest possible aggregation, which ignores binding entropy entirely. The Phe/Tyr comparison adds mechanistic support, as TPM score differences tracked solvation-inclusive but not gas-phase quantum-mechanical interaction energies, consistent with the representation reflecting the solvated environments from which it was learned. Together, these analyses show that a single native-structure representation captures both where an interface is sensitive to sequence perturbation and which alternative residues are locally preferred, the two quantities most directly constrained by local interface context.

These results bear out the premise with which we began. Repeated observation of which interaction motifs occur in which local structural environments is sufficient to recover information about experimentally measured mutation sensitivity and substitution preference without quantitative affinity labels. This offers a complementary route to direct ΔΔ*G* learning, for which experimental training data remain sparse relative to the diversity of possible protein interfaces. Rather than learning free-energy changes directly, TPMs learn recurring structural signatures of compatible molecular environments, from which energetically relevant information emerges.

The scope of this information is nevertheless set by the representation. The present study evaluates TPMs from individual experimentally resolved structures and therefore does not explicitly model conformational rearrangement, induced fit or configurational entropy, effects that are particularly important for absolute binding free energies and that may partly explain why the whole-interface affinity association remains modest even where local mutation effects are recoverable. The formulation itself is not restricted to static structures, and applying the same mapping across conformations from molecular dynamics simulation or mode-guided sampling could expose how preference fields change with protein motion, as illustrated in Extended Data Video 1 (Methods 3.18). Therefore, such effects are better represented by the input than excluded by the method. TPMs also inherit uncertainty from the structural data used for training and prediction, and errors in local geometry, side-chain or protonation states such as ambiguous histidine tautomers, crystal contacts or biological-assembly assignment can propagate to the inferred interaction environments.

A second boundary follows from the deliberately target-centred representation. TPMs describe where a receptor environment favours a defined interaction motif, but they do not encode whether those locally preferred motifs can be connected into a chemically and conformationally feasible binder, so ligand connectivity and global realizability must be supplied by another component when moving from interface analysis to design. Training coverage imposes a further constraint, since interaction modes not represented in the structural record cannot be expected to emerge reliably, as illustrated by the poorer transfer of cysteine-associated motifs and the near-absence of interfacial disulfide geometries in the antibody–antigen training set. More generally, structural databases are biased toward well-studied protein classes, experimentally tractable complexes and strongly interacting systems, and these biases will shape the interaction environments available for learning.

A final set of limitations concerns the current readout of the learned fields rather than the fields themselves. Substitution ranking was strongest at geometrically embedded sites and deteriorated at solvent-exposed or loosely packed positions, and FoldX and Flex ΔΔ*G* showed the same dependence, suggesting that local geometric embedding is a general axis of mutation-effect predictability rather than a limitation specific to TPMs. The fixed-frame rotamer and pose search adds a further approximation because candidate substitutions are evaluated without side-chain or local backbone relaxation, and the 1PPF Y20 example shows that the underlying field can retain informative geometry even where the scalar score from the placed rotamer is imperfect. The present fit function should therefore be regarded as a simple readout of the information contained in the fields rather than an optimized mutation-energy estimator. That this readout nevertheless performs comparably to dedicated ΔΔ*G* methods indicates that binding-relevant signal is present in the learned fields, and that further gains may be possible by coupling them to conformationally adaptive placement or relaxation.

The main opportunity opened by TPMs is their combination of transferability, energetic relevance and spatial interpretability. Because predictions remain resolved as fields for defined interaction motifs, the same representation can localize regions sensitive to sequence perturbation, prioritize alternative substitutions and expose the interaction geometry underlying those preferences, without task-specific training. For interface optimization, this could provide a rapid structural filter alongside physics-based mutation scoring, particularly when many candidate substitutions must be evaluated from a native complex. For generative design, the separation between local preference and ligand connectivity becomes an opportunity, as TPM fields could supply explicit chemical conditioning to a generative model that provides the globally connected binder, concentrating sampling in regions compatible with molecular recognition while preserving alternative solutions through overlapping fields for chemically related motifs. The present results suggest that structural databases encode more than examples of successful binding geometries. Repeated observation of local molecular environments can itself provide supervision from which transferable, energetically relevant interaction preferences emerge. This offers a route to learning protein-interface energetics from the much larger structural record even where quantitative affinity and mutation measurements remain sparse. More generally, local preference fields provide a bridge between structural machine learning and molecular interpretation: they retain the flexibility of data-driven prediction while remaining anchored in atomic interaction geometry and experimentally observed molecular recognition.

## 3 Online Methods

### 3.1 Shard definition and interface decomposition

We represent protein interfaces as collections of sparse local chemical features. Each interaction motif, or shard, is represented by one to three anchor atoms that define reference points for local target environments. The spatial arrangement of an anchor’s preferred local environment is learned by a separate classifier, so that a shard with multiple anchors is described jointly by several anchor classes. The model uses 36 anchor classes spanning backbone-associated, charged, polar, aromatic and hydrophobic features (Extended Data Table 2). For readability, anchor classes are referred to throughout the manuscript by residue–atom labels such as Lys-NZ, Arg-CZ and Tyr-OH. The anchor scheme is a sparse representation rather than an exhaustive atom-wise decomposition of each residue. Anchor positions were selected during model development to provide spatially separated reference points that were expected to yield localized, orientation-sensitive preference signals, and were then kept fixed across training and all downstream analyses.

Three residue-independent backbone classes, BB-C*β*, BB-O and BB-H, represent the C*β* position, backbone carbonyl oxygen and backbone amide hydrogen, respectively. C*β* and the amide hydrogen were chosen instead of C*α* and backbone nitrogen because their positions provide greater spatial separation from the backbone and were expected to yield more discriminative spatial signals. The amide hydrogen was placed from the local covalent geometry during structure preparation. Glycine, which lacks a C*β* atom, is represented separately by the Gly-C*α* class.

A single chemical group is therefore represented by one or more anchor classes at distinct positions. For example, the arginine guanidinium is represented jointly by the Arg-CZ and Arg-NH anchor classes, and the histidine imidazole by the His-CG, His-ND1 and His-NE2 classes. Where chemically equivalent atoms occupy interchangeable positions, they are grouped within the same anchor class, so that Arg-NH includes both NH1 and NH2, with analogous grouping for symmetric carboxylate and aromatic atoms.

Because each anchor independently defines the origin of a local target environment, the spatial arrangement of multiple anchor predictions encodes the geometry and orientation of an interaction (Fig. 1a,c). For antibody–antigen complexes, interfaces were decomposed in both directions, with each partner contributing anchors in turn while atoms of the opposing partner defined the corresponding target environments.

### 3.2 Datasets and structural preprocessing

Training structures were obtained from SAbDab antibody-antigen complexes [19] in Chothia numbering. Antibody heavy and light chains and their corresponding antigen chains were identified from the SAbDab REMARK annotations. Repeated identical pairing annotations were ignored, and antibody chains associated with the same antigen were combined where appropriate.

Structures were parsed from ATOM records only. Waters, ions and other HETATM records were excluded. Deposited hydrogen atoms were removed and a single backbone amide hydrogen was reconstructed for each non-proline residue by placing it on the backbone nitrogen anti-parallel to the preceding carbonyl bond. Alternate conformations were resolved by retaining the highest-occupancy conformer. Histidine protonation and tautomer states are not consistently resolved in deposited protein structures and generally require structure-specific inference from the local environment. Histidine protonation variants were therefore canonicalized to a single residue identity, and the two imidazole nitrogens were assigned the same atom class rather than distinguished by protonation state or tautomer. Structures were excluded if the annotated interaction partners shared chains, if required chains were absent from the coordinate records, if an unknown residue (UNK) occurred in a relevant chain, if relevant chains contained only C*α* atoms, or if three or more backbone atom types (N, C*α*, C or O) were missing. A total of 6291 antibody–antigen structures passed preprocessing and were carried forward for environment extraction.

For each interface-extraction direction, a permissive distance pre-filter was applied to limit computation. Candidate shard-bearing atoms were retained if they lay within 13 Å of any atom on the opposing interaction partner, and target atoms were retained if they lay within 35 Å of any retained shard-bearing atom. These cutoffs do not define the local interaction environment. The operative neighbourhood and subsequent validity filtering are defined during environment encoding below.

### 3.3 Encoding of local target environments

The representation uses four related terms. A shard is a rigid chemical group. Each shard is located and oriented by one or more shard anchors, the atoms that mark its defining positions. The set of equivalent anchor atoms modelled by a single classifier is an shard anchor (36 in total, Extended Data Table 2). Separately, each surrounding target atom is labelled by an atom class, a chemical type used to describe the local environment (Extended Data Table 3). Anchor classes thus index the models, whereas atom classes describe the environment those models read. For each shard anchor, atoms of the opposing interaction partner were represented by their chemical atom class and their position relative to the anchor.

Spatial information was represented by displacement vectors (Δ*x,* Δ*y,* Δ*z*) from the shard anchor to each target atom. Coordinates were expressed in the translated global coordinate frame of the input structure. No rotation into a shard-specific local frame was applied. Target atoms farther than 40 Å from the anchor were excluded.

Encoded environments retained up to the 100 nearest target atoms. Neighbours were ordered by distance to the anchor, which allows for easier inspection against the corresponding structures. Shorter environments were zero-padded, using a reserved atom class (class 0) and a zero displacement vector to distinguish padding from real atoms. The models themselves use only the 25 nearest neighbours from these encoded environments (Section 3.6).

To exclude sparse or poorly defined local environments, we applied a fixed validity filter to these 25 nearest neighbours. An environment was retained only if at least five neighbours lay within 8 Å of the anchor, no more than three lay beyond 13 Å, and at least two distinct target-atom classes were present. This filter defines the local environments used for subsequent model training and evaluation.

### 3.4 Positive and negative training examples

For each anchor class, positive examples were the native local environments centred on anchors of that class in antibody-antigen interfaces. Negative examples were generated from two complementary sources and combined in equal proportion during training.

The first source comprised random surface environments, sampled from solvent-accessible regions around the protein and encoded identically to native environments. Candidate positions were generated on a 0.5 Å grid around the molecular surface, excluding points within 2 Å of any protein atom and progressively down-weighting positions farther from the surface. The number of sampled positions was proportional to solvent-accessible surface area, producing a distribution of non-native local environments surrounding the protein surface (Extended Data Fig. A1).

The second source comprised cross-shard negatives, native environments centred on anchors belonging to other shard classes. Distinct anchors of the same chemical group were retained as negatives for one another. For example, the second carbox-amide anchor contributes negatives to the classifier trained for the first. This requires each classifier to distinguish the local environment associated with its specific anchor position rather than only the broader chemical identity of the group.

Negative examples were resampled at each training epoch rather than drawn from a fixed negative set. Random surface environments were used only during training. All reported shard-discrimination AUCs were calculated using native cross-shard environments as negatives, thereby testing discrimination between experimentally observed environments of different shard classes rather than between native interfaces and arbitrary surface positions.

### 3.5 Training, validation and test splits

Redundancy between training, validation and test sets was controlled at the antibody complementarity-determining-region (CDR) sequence level. For each complex, the six CDR loops (H1-H3 and L1-L3) were extracted according to Chothia numbering and concatenated into a single sequence. Conserved framework regions were excluded from clustering. For complexes containing multiple antibody pairs against a shared antigen, the corresponding CDR sequences were concatenated consistently with the interface-extraction chain assignment. CDR sequences were clustered with CD-HIT at 70% sequence identity, and complete clusters were assigned to training, validation or test sets in an approximately 80/10/10 split, preventing CDR-similar complexes from spanning data partitions.

Complexes containing SARS-CoV-2 spike protein were assigned to the test set at cluster level, ensuring that the SARS-CoV-2 complexes used for downstream evaluation and structural examples were excluded from training. Model selection, including checkpoint selection and early stopping, used the validation set only. The test set was not used during training or model selection. CDR-based clustering provides a stringent sequence-level separation of antibody interfaces across data partitions. As an additional safeguard against residual similarity within the antibody-antigen domain, we evaluated the models on independent peptide-protein and non-antibody protein-protein complexes that lie outside the training interaction class.

### 3.6 Model architecture and training

Each of the 36 anchor classes was modelled independently using a binary classifier with a shared architecture and separately trained weights. For a given anchor, the model predicts whether an encoded local environment corresponds to a native environment of that shard class. Training batches were balanced between positive and negative examples, with negatives sampled as described above.

Each environment was represented by the 25 nearest target atoms through two input channels. First, their displacement vectors (Δ*x,* Δ*y,* Δ*z*) relative to the candidate shard anchor and second their corresponding atom-class tokens. Displacement vectors were normalized using a feature-wise normalization layer adapted on 20000 sampled training environments. Atom-class tokens were mapped to learned 32-dimensional embeddings, and the resulting atom and coordinate representations were concatenated for each neighbour, giving a 35-dimensional per-atom representation.

These representations were processed by a Perceiver classifier [26, 27]. A trainable array of 32 latent vectors cross-attends to the 25 per-atom representations, followed by a self-attention transformer over the latent array, with this cross-attention and transformer step applied for 2 iterations. The transformer used 4 self-attention blocks with 4 attention heads each and feed-forward sub-networks of hidden sizes [128, 35]. The final latent representation was reduced by global average pooling and passed to a classification head of hidden sizes [128, 64, 1] with a sigmoid output. To encourage rotational invariance in a model that operates directly on Cartesian displacement vectors, each local environment was augmented by random three-dimensional rotations (n=12) during training, so that the classifier learns from the relative geometry of the environment rather than any absolute orientation. Models were optimized using binary cross-entropy loss and Adam with a learning rate of 10*^−^*^4^ and batch size 24. Training proceeded for up to 15 epochs with early stopping after five epochs without improvement in validation loss, and the checkpoint with the lowest validation loss was retained. Negative examples were resampled at every epoch as described above. Random seeds for dataset partitioning and model training were fixed for reproducibility.

### 3.7 Target preference map generation

A target preference map (TPM) is obtained by evaluating a trained classifier over a regular three-dimensional grid surrounding a target protein. At each grid point, the classifier estimates the compatibility of the local protein environment with placement of the corresponding shard anchor at that position. Target structures were processed using the same atom-class representation as during training and restricted to the selected receptor chains. Depending on the analysis, this region encompassed a known interaction site, the complete target surface or a local residue-centred neighbourhood. Each voxel centre was treated as a candidate shard-anchor position. The 25 nearest target atoms were retrieved and represented by their atom classes and displacement vectors relative to the candidate position, using the same encoding as during training. The corresponding shard classifier was then evaluated to obtain a compatibility for each voxel, yielding a three-dimensional scalar field. TPMs for different shard anchor classes were calculated independently on the same coordinate grid. For chemical groups represented by multiple classes, the relative spatial arrangement of their maps therefore specifies preferred interaction geometry without requiring an explicit orientation variable. Unless otherwise stated for a specific downstream analysis, TPM values correspond to the raw model output. TPMs were stored as MRC volumetric maps with the grid origin and voxel spacing retained in the file header, and as numerical arrays for quantitative analysis.

### 3.8 Held-out antibody–antigen and peptide–protein evaluation sets

Generalization within the antibody–antigen domain was evaluated on the held-out test partition described above, which was separated from the training data at the level of 70%-identity antibody CDR-sequence clusters.

To test transfer beyond antibody–antigen recognition, we constructed an independent peptide–protein benchmark from the protein–protein subset of PDBbind. For each complex, chain lengths were determined from standard amino-acid residues in the first structural model. Complexes were initially retained when they contained at least one peptide-like chain of at most 30 residues and a distinct protein chain of at least 80 residues. When multiple chains satisfied these criteria, the shortest qualifying chain was assigned as the peptide and the longest distinct qualifying chain as the protein. This initial screen yielded 579 candidate complexes.

To obtain an unambiguous two-partner benchmark, candidates were further restricted to structures containing exactly two protein chains. This removed multichain complexes in which assignment of the peptide–protein interface could be ambiguous and reduced the benchmark from 579 candidates to 185 complexes. The peptide chain was treated as the shard-bearing interaction partner and the longer protein chain as the target environment. Only this interaction direction was evaluated because the short peptide does not provide an extended protein-like target environment for the reverse direction.

Local environments were generated using the same structural preprocessing, atom representation and validity criteria as for the antibody–antigen data. None of the 185 peptide–protein complexes shared a PDB identifier with the SAbDab structures used for model development, and the peptide–protein set was not used for training, hyperparameter selection or model selection.

### 3.9 Shard-level discrimination analysis

Shard-level discrimination was evaluated separately on the held-out antibody–antigen and independent peptide-protein sets. For each classifier, positive examples were native environments belonging to that shard class, whereas negative examples comprised native environments belonging to all other shard anchor classes in the same evaluation set. Thus, negatives included chemically related and alternative anchors of the same functional group, making the task stricter than discrimination between broad chemical classes. Random surface environments used during training were not included in evaluation.

Each classifier was applied to all corresponding positive and negative environments, and discrimination was quantified by the area under the receiver operating characteristic curve (ROC-AUC). AUCs were calculated independently for each of the 36 shard anchor classes from the pooled environments in each evaluation set. The same scored environments were subsequently used for the pairwise shard-discrimination analysis described below.

### 3.10 Environment enclosure analysis

To quantify how completely the target protein surrounds a shard anchor, we calculated an enclosing solid-angle measure for each valid local environment. The displacement vectors from the anchor to the 25 neighbouring target atoms used for prediction were converted to unit direction vectors *v*^*_i_*. We then determined the axis *u*^ of the smallest cone containing all neighbour directions by maximizing the smallest projection min*_i_*(*û_i_ · u*^). The resulting cone half-angle is

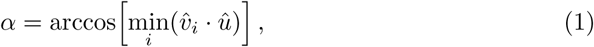

and enclosure was expressed as the fraction of the full sphere subtended by this cone,

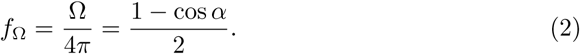

Small *f*_Ω_ values indicate that neighbouring target atoms lie predominantly on one side of the anchor, as expected for an exposed surface position, whereas larger values indicate that the anchor is increasingly surrounded by the target protein, as in a groove or pocket. Padding entries were excluded from the calculation.

For the comparison in Fig. 2b, *f*_Ω_ was calculated independently for the valid shard environments in the held-out antibody–antigen and peptide–protein evaluation sets, and the resulting environment-level distributions were compared.

### 3.11 Pairwise shard discrimination

The aggregate shard-level AUC described above asks whether a shard anchor classifier can distinguish its own native environments from the pooled native environments of all other shard anchor classes. To resolve which specific shard classes occupy distinct or overlapping local environments, we repeated this analysis pairwise.

For every ordered pair of shard anchor classes (*i, j*), the classifier trained for shard *i* was evaluated on two sets of experimentally observed environments: native environments of shard anchor *i*, treated as positives, and native environments of shard anchor *j*, treated as negatives. The corresponding matrix entry was defined as

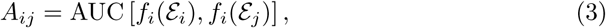

where *f_i_* denotes the classifier for shard anchor *i* and *E_i_* and *E_j_* denote the native environments associated with shard anchors *i* and *j*, respectively. Equivalently, *A_ij_*measures the probability that the shard anchor-*i* classifier assigns a higher score to a randomly selected native environment of shard anchor *i* than to a randomly selected native environment of shard anchor *j*. The diagonal entries *A_ii_* are undefined because the positive and comparison classes are identical.

Thus, each row of the matrix asks a different question: given the environmental preferences learned for the row shard anchor, how distinguishable are the native environments of each column shard anchor? Values approaching 1 indicate that the row classifier strongly separates its own native environments from those of the column shard anchors. Values near 0.5 indicate strongly overlapping score distributions, meaning that the row model does not distinguish the two environmental classes under its learned representation. Values below 0.5 indicate that environments of the column shard anchor tend, on average, to receive higher scores from the row classifier than the row shard anchor’s own environments. Reduced off-diagonal AUCs are therefore not necessarily classification failures. Particularly for chemically related shard anchors, they can reflect genuine overlap in the local structural environments compatible with the two interaction motifs.

The matrix is asymmetric by construction. *A_ij_* is calculated with the shard anchor-*i* classifier, whereas *A_ji_* is calculated with the independently trained shard anchor-*j* classifier. The two quantities therefore need not be equal. For example, if environments recognized by shard anchor*i* form a relatively restricted subset of those tolerated by shard anchor *j*, the shard anchor-*i* model may distinguish *i* from *j* more strongly than the shard anchor-*j* model distinguishes *j* from *i*. This directional information allows chemically related motifs to share compatible environments without requiring their learned preferences to be identical.

Pairwise matrices were calculated independently for the held.-out antibody-antigen and peptide-protein evaluation sets using the same prediction scores as the aggregate shard-level analysis. For each row shard anchor, the aggregate AUC reported above is therefore the corresponding one-versus-rest calculation in which the negative set is the union of all shard anchor classes *j ̸*= *i*, whereas the pairwise matrix resolves this pooled comparison into its individual shard-shard anchor components. No random-surface environments were included in the pairwise analysis.

### 3.12 Phe/Tyr QM substitution benchmark

To test whether TPM score differences carry graded energetic information beyond categorical discrimination, we compared them with quantum-mechanical (QM) substitution energetics for matched phenylalanine/tyrosine substitutions. Phe and Tyr differ by a single hydroxyl group, providing a defined chemical perturbation with otherwise closely matched local geometry. The analysis was performed exclusively on the independent peptide–protein evaluation set (Section 3.8), which was not used for model training or selection.

Substitution sites comprised native Phe and Tyr residues at peptide–protein interfaces, each analysed in its native and substituted identity. For TPM scoring, substitutions were generated by adding or removing the hydroxyl oxygen while retaining the shared aromatic-ring coordinates. No additional modelling of hydroxyl orientation was required because TPM scoring uses the oxygen position directly. For the QM calculation, both orientations of the Tyr phenol were constructed and the orientation with the lower interaction energy was retained.

QM interaction energies were calculated with In-Pocket-Analysis (IPA) using GFN2-xTB and an energy-decomposition scheme over the complex and its separated partners. Local QM regions were selected by IPA using a 10 Å residue-inclusion radius around the substitution site. Energies were evaluated with implicit water solvation using the generalized Born model and, as a control, in the gas phase. In both cases the IPA bias-potential parameters were fixed at *α* = 1.0 and *κ* = 0.025.

For each site, differences were defined using a native-minus-mutant convention. ΔTPM was the difference in summed TPM score between the native and substituted residue, and Δ*E*_QM_ the corresponding difference in IPA interaction energy. Their association was quantified by Spearman rank correlation with bootstrap 95% confidence intervals, using the solvated energies as the primary comparison and gas-phase energies as a control. A total of 182 sites yielded defined ΔTPM and Δ*E*_QM_ values and entered the analysis.

### 3.13 SKEMPI mutation benchmark and analysis subsets

#### Source and primary filtering

The mutation benchmark was derived from SKEMPI 2.0. We retained only single-point substitutions at binding-interface positions, defined as core, support or rim residues (COR, SUP and RIM) according to the Levy interface classification [29]. Interior and non-interface surface positions were excluded. Binding free-energy changes were recalculated directly from the reported wild-type and mutant dissociation constants rather than taken from a pre-derived ΔΔ*G* field:

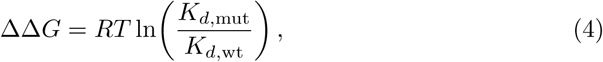

using the temperature reported for each measurement. Only entries with finite, positive *K_d_* values for both states were retained. Positive ΔΔ*G* denotes a destabilizing substitution. Multiple measurements of the same substitution were averaged.

Because TPM was trained exclusively on antibody-antigen complexes, the benchmark was restricted to interactions outside this training class. Antibody-antigen (AB/AG) and T-cell-receptor/peptide-MHC (TCR/pMHC) complexes were excluded, leaving protease-inhibitor and other non-antibody protein-protein interactions. This exclusion was important for leakage-free evaluation. 41 of the 45 filtered AB/AG complexes shared a PDB identifier with structures used for TPM training, whereas none of the retained complexes did.

#### Full mutation benchmark

The resulting parent benchmark comprised 217 complexes, 2833 interface substitutions and 1323 unique mutated interface positions. This set underlies the mutation-sensitive-position analysis (Fig. 3).

#### Mutation-sensitive positions

Each of the 1323 positions was assigned a binary label from its experimental mutation series. A position was defined as mutation-sensitive if at least one measured substitution satisfied |ΔΔ*G| ≥* 1.0 kcal mol*^−^*^1^, and mutation-tolerant otherwise. TPM scoring for this analysis used the wild-type structure alone, without modelling mutant residues.

#### Ranking benchmark

Substitution ranking required sufficient experimental coverage and non-degenerate TPM scores. Pockets were retained when fewer than half of their measured substitutions received a TPM score of exactly zero and when at least 15 substitutions had been measured. These criteria yielded 33 pockets for the per-pocket ranking analysis (Fig. 4).

#### Specialized nested subsets

Two subsets of these 33 pockets were defined for additional analyses. The sign-discriminating subset contained at least one substitution with ΔΔ*G < −*0.5 and one with ΔΔ*G >* +0.5 kcal mol*^−^*^1^, yielding 15 pockets.

The embedded subset contained pockets with a side-chain embeddedness of *≥* 0.04, where side-chain embeddedness is the occluded solid-angle fraction of the native side-chain anchor atoms (the atoms scored by TPM), computed as described below, yielding 21 pockets.

#### Side-chain embeddedness

Side-chain embeddedness quantifies how strongly the native side-chain anchor atoms of a mutated position are enclosed by the opposing interaction partner. For each pocket, the reference atoms were the native side-chain anchor atoms of that residue, that is, the functional-group atoms scored by TPM rather than the full residue including backbone. Referencing the metric to these anchor atoms measures burial of the atoms TPM actually scores and avoids the backbone contribution that dominates an all-atom contact measure at functionally exposed side chains.

The sphere of directions around these reference atoms was discretized into a grid of approximately equal-area cells (18 latitude, 36 longitude cells. 648 cells in total), and each partner heavy atom within an 8 Å shell was assigned to the cell containing its direction. The occluded solid-angle fraction is the fraction of the 648 cells occupied by at least one partner atom, normalized over the full sphere (4*π*). Because partner atoms occupy only a patch on the interface-facing side of a residue, even a well-enclosed side chain covers only a small fraction of the full sphere. The metric is therefore a relative measure of directional coverage rather than an absolute ”fraction enclosed”. The absolute values and the *≥* 0.04 threshold are specific to this bin resolution and shell radius. Pockets with an occluded solid-angle fraction *≥* 0.04 were assigned to the embedded subset. The per-pocket zoom-in views in Extended Data (Figs. A5–A25), together with the summary scatter (Extended Data Fig. A4), illustrate this metric visually, spanning embedded to peripheral pockets.

### 3.14 Native TPM scoring of mutation-sensitive interface positions

We asked whether a TPM score computed from the wild-type complex alone identifies interface positions that are sensitive to mutation. Only the native residue was scored. No mutant structure, conformational search or structural relaxation was used.

#### Native per-residue TPM score

For each mutated interface position, the trained shard anchor classifiers of the native residue were evaluated at the experimentally resolved anchor coordinates. For every residue-specific shard anchor, the corresponding classifier was applied to the local environment at that position using the same representation as during training and TPM generation (Sections 3.3 and 3.7). Scores were sampled directly at the native coordinates without searching nearby positions, yielding one compatibility score per valid shard anchor.

The per-residue score was defined as the mean compatibility score over the residue-specific shard anchors. The generic BB-C*β*, BB-O and BB-H classes were excluded because they are shared across residue identities and therefore do not encode residue-specific chemistry. Their inclusion would also duplicate anchor positions represented by residue-specific shards in several residue classes. Residue-specific classes such as Gly-CA and Ala-C*β* were retained. Invalid shard environments were omitted from the mean, and a position was scored when at least one residue-specific anchor was valid. No residue-type normalization or additional correction for the number of contributing anchors was applied.

#### Mutation-sensitive labels and discrimination

Each of the 1323 SKEMPI interface positions was assigned the binary label defined in Section 3.13. A position was considered mutation-sensitive when at least one measured substitution satisfied |ΔΔ*G| ≥* 1.0 kcal mol*^−^*^1^ and mutation-tolerant otherwise. Because the TPM score is calculated once from the wild-type residue, the analysis was performed at the level of unique positions rather than individual substitutions. Discrimination between mutation-sensitive and mutation-tolerant positions was quantified by ROC-AUC using the native per-residue TPM score as the ranking variable and the mutation-sensitive label as the positive class.

### 3.15 TPM scoring of interface substitutions

To rank alternative substitutions at a fixed interface position, candidate amino acids were scored against target preference fields generated from the wild-type complex. The resulting fit score measures geometric and chemical compatibility with the fixed local environment. It is not a predicted binding free-energy change and uses no experimental affinity data.

#### Fixed structural environment and field masking

The wild-type complex provided the fixed structural environment, and TPM fields were computed once on the wild-type receptor (Section 3.7). Candidate side chains were fitted on the native backbone without backbone remodelling, receptor relaxation or mutant-complex minimization, and TPM fields were not recomputed for individual candidates. Before scoring, fields were Gaussian-smoothed and masked at voxels occupied by protein atoms of the ligand. At the mutated position, only backbone atoms contributed to this self-chain mask, leaving the native side-chain volume accessible to candidate residues. C*β*-centred shards used a reduced exclusion radius.

#### Candidate construction and torsion search

Each interface position was represented in a local frame defined by its native backbone, with C*α* as the origin, the C*α →*C*β* direction as the principal axis and C*α →*N fixing the handedness. Candidate side-chain functional-group atoms were generated from fixed internal-coordinate templates in this frame, preserving the native N, C*α* and C*β* geometry. Only side-chain torsions were varied. Bond lengths, bond angles and backbone coordinates remained fixed. Gly, Pro, Cys and Met were excluded from candidate substitution scoring.

All side-chain *χ* angles were searched exhaustively over [*−*180 deg, +180 deg) in 30 deg increments. All combinations were enumerated by Cartesian product, giving 12*^n^* poses for a residue with *n* rotatable torsions (up to 12^4^ = 20736 poses for Arg). Each rotation affected only atoms distal to the corresponding bond. No additional rigid-body translation or rotation of the side chain was applied.

#### Per-pose and substitution scoring

Candidate residues were represented by residue-specific shard anchors. The generic BB-C*β*, BB-O and BB-H classes were excluded. Almost all shard anchors were treated as primary. The only secondary anchors were the distal indole carbon of tryptophan (CZ3) and the tyrosine hydroxyl (OH), which are rotationally labile and least reliable for fixing the rigid side-chain pose. For a primary anchor *a* at posed position **x***_a_*, its shard field was searched within 1.0 Å and field values were penalized according to their displacement from the posed anchor using a Gaussian with *σ* = 0.5 Å. The anchor score was the maximum distance-penalized field overlap within this neighbourhood, and the primary pose score was the mean across all primary anchors.

The pose with the highest primary score was retained. Secondary anchors were evaluated only for this pose and contributed a bounded bonus using a field threshold of 0.60, a search radius of 3.0 Å and weight 0.20. Each anchor was evaluated against its own shard-specific field. Fields belonging to different anchors were not merged.

Primary scores were subsequently rescaled using fixed chemistry-class factors (directional, *×*0.80; hydrophobic and aromatic, *×*1.00), after which the secondary-anchor bonus was added. The resulting value is the raw TPM fit score for the candidate amino acid, with higher values indicating greater compatibility with the fixed interface environment. Scores are absolute candidate-fit scores and are not referenced to the wild-type residue.

#### Fit parameters and applicability

The local search tolerances, Gaussian penalty, chemistry scaling and secondary-anchor contribution provide a simple allowance for small placement errors in the otherwise rigid fitting procedure. These parameters were set once by inspection of one TPM field and then held fixed for all benchmark positions. Experimental ΔΔ*G* values were not used to select or optimize them.

A candidate received a score of zero when none of its primary anchors could be placed in admissible, non-masked field density at any searched pose. Pockets dominated by zero-scoring candidates were excluded according to the applicability criterion defined in Section 3.13. Raw TPM scores are defined such that higher values indicate greater compatibility. Neither candidate generation, pose selection nor TPM scoring used experimental mutation labels.

### 3.16 Mutation-pocket ranking analysis

Substitution ranking was evaluated independently within each of the 33 SKEMPI pockets defined in Section 3.13, and compared against two structure-based baselines, FoldX and Flex ddG. Precomputed FoldX and Flex ddG predictions for SKEMPI2.0 were obtained from the baseline data released by Deng et al. [30] and were not recomputed in this study. To ensure a strictly matched comparison, the three methods were evaluated on an identical set of substitutions within each pocket. For every pocket, only substitutions for which a TPM fit score, a FoldX prediction and a Flex ddG prediction were all available were retained, and all three methods were scored on this common set. Substitutions lacking a prediction from any method were excluded from that pocket for all methods, so that no method was evaluated on substitutions unavailable to another.

For each pocket, the predicted scores of the retained substitutions were correlated with their measured ΔΔ*G* values by Spearman rank correlation. All predictors were expressed in the experimental ΔΔ*G* convention, in which a higher value denotes a more destabilizing, less favourable substitution. The raw TPM fit score increases with predicted compatibility and therefore runs opposite to ΔΔ*G*, so it was sign-inverted to *−*TPM. FoldX and Flex ddG already report ΔΔ*G* in this convention and were used without modification. Under this common convention, a positive Spearman coefficient indicates that a method ranks the substitutions in the correct experimental order within the pocket, and *−*TPM is the quantity plotted against measured ΔΔ*G* in Fig. 4d-g.

Each pocket contributed a single Spearman coefficient per method, irrespective of the number of substitutions retained at that pocket. Aggregate performance was summarized by the unweighted mean of the per-pocket coefficients, so that each pocket contributed equally regardless of how densely it had been mutagenized. Uncertainty in each aggregate mean was estimated by bootstrap resampling of pockets. Pockets were resampled with replacement 10000 times, the unweighted mean per-pocket correlation was recomputed for each resample, and the 2.5th and 97.5th percentiles of the resulting distribution defined the reported 95% confidence interval. Resampling was performed over pockets rather than substitutions, consistent with the pocket-level unit of analysis.

The same procedure was applied to the two nested subsets defined in Section 3.13. The sign-discriminating subset, comprising pockets that contain both experimentally stabilizing and destabilizing substitutions, assessed whether a method separates favourable from unfavourable substitutions at a single position. The embedded subset (side-chain embeddedness *≥* 0.04) assessed ranking at geometrically constrained, more embedded positions. The identical aggregation and bootstrap procedure was applied to each subset independently. The common-substitution restriction was applied within every pocket of every subset, so that all reported comparisons rest on the same substitutions across the three methods.

This analysis measures ranking within pockets and does not test whether TPM fit scores are comparable across pockets. The raw fit score is not calibrated to an absolute binding free-energy scale, so no pooled, cross-pocket correlation was computed. Each pocket was assessed on its own substitutions only.

### 3.17 Global interface-affinity analysis

To test whether TPM compatibility scores, aggregated over an entire interface, track absolute binding affinity, we defined a single unweighted interface-level score and correlated it with measured affinity across two independent interface sets. This analysis deliberately uses no fitting of any kind. No per-family weights, no interface-area normalization and no exclusion of low-performing shard anchor classes. It therefore tests the raw aggregate signal rather than any tuned combination of it.

#### Interface aggregate

For each interface, every anchor of the interacting partner was scored by its corresponding shard anchor classifier as in TPM generation (Section 3.7), yielding one compatibility score per anchor. Anchors were retained if their compatibility score was at least 0.5. Retained anchors were grouped by residue, the mean compatibility score was taken within each residue, and these per-residue means were summed to give a single score per interface. Aggregating by residue mean before summing prevents residues bearing many anchors from dominating the score purely through anchor count, while the sum over residues still increases with the number of well-satisfied interface residues. No normalization by interface area or residue count was applied.

#### Affinity data

Two interface sets were analysed. The first comprised the SARS-CoV-2 antibody/nanobody and ACE2–RBD complexes used elsewhere in this study, with binding affinities expressed as pK_d_ = *−* log_10_(*K_d_/*1 M). The second comprised the PDBbind peptide–protein complexes of the independent benchmark (Section 3.8). For this set, affinities were taken from the PDBbind index. The reported binding constant (*K_d_*, *K_i_*or IC_50_) was converted to molar units and expressed as *−* log_10_(affinity*/*1 M), so that higher values denote tighter binding on both axes. Because the index reports a mixture of constant types, the peptide axis is a pooled affinity measure rather than *K_d_* alone. *K_i_* and IC_50_ approximate *K_d_* under standard assumptions and were treated interchangeably for this analysis, which asks only whether the interface aggregate tracks affinity at all. Only measurements reported as exact (= relation) were retained.

### 3.18 Animation of preference fields

To illustrate that preference fields are not restricted to a single fixed geometry, we generated a conformational trajectory of hen egg-white lysozyme (PDB 1LYZ) and recomputed TPMs along it. Non-protein atoms were removed and missing atoms and hydrogens were added with PDBFixer at pH 7.0. A conformational trajectory was then generated with the ANMD protocol in ProDy [31], which displaces the structure along an anisotropic network model (ANM) mode and relieves the resulting distortions by all-atom energy minimization at each step. We followed the lowest-frequency non-trivial ANM mode to a maximum C*α* RMSD of 2 Å in 10 steps, yielding 11 conformers spanning the motion (default ANM parameters: 15 Å cutoff, uniform spring constants). For each conformer, TPMs were computed for the Lys-NZ, Asp/Glu-C and Asn/Gln-C anchor classes (Extended Data Table 2) using the same procedure as for static structures (Methods 3.7). The resulting fields were rendered per conformer to produce Extended Data Video 1, in which the preference fields redistribute as the backbone moves along the mode.

## Declarations

### Ethics approval and consent to participate

Not applicable.

### Consent for publication

Not applicable.

### Data availability

The datasets analyzed in this study are derived from publicly available resources as described in the Methods. Processed datasets and benchmark data will be made available upon publication of the peer-reviewed article.

### Code availability

Code for model training, TPM generation and analysis will be made available upon publication of the peer-reviewed article.

### Author contribution

G.M.P. and M.K. designed the study. M.K. implemented the software, performed the analyses and wrote the manuscript. G.M.P. conceived and supervised the project and acquired funding. F.M. contributed to method development and analysis. G.M.P., F.M., P.G., S.M. and T.S. reviewed and edited the manuscript.

