## Extended Data for "Local structural preference maps encode transferable protein-interface energetics"

### Appendix A  Extended Data

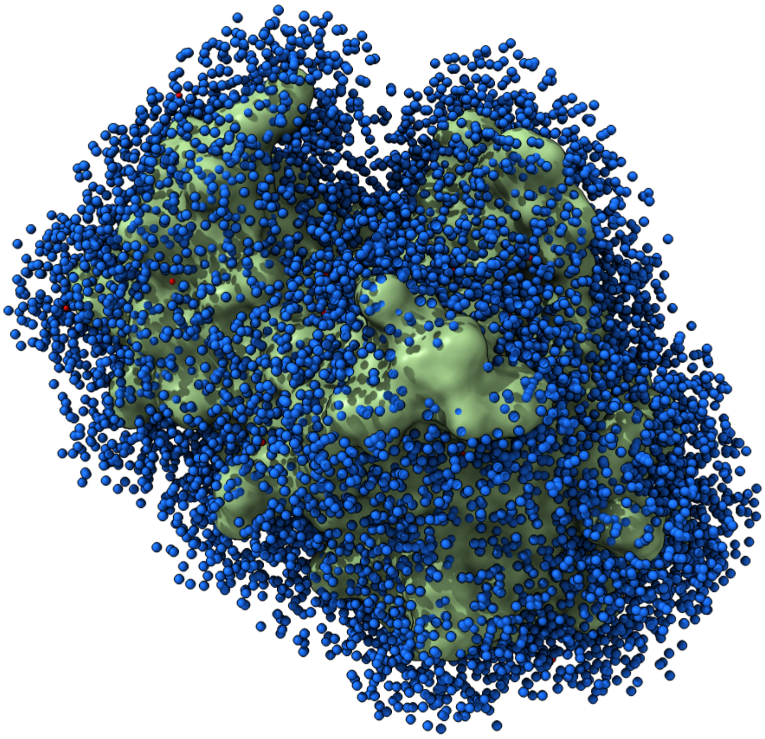

**Fig. A1 Spatial distribution of sampled negative points around a target protein.** A representative structure shown in surface representation, with sampled negative-environment centres rendered as spheres. Negative points are drawn from the solvent-accessible region surrounding the protein. Candidate positions were generated on a voxel grid (0.5 Å spacing, 4 Å padding) and sampled with a distance-dependent weight that excludes positions within 2 Å of any atom and decreases linearly with distance thereafter, yielding an effective sampling shell of approximately 2 Å to 14.5 Å from the surface. The number of points scales with solvent-accessible surface area ($n = n_0(A/A_0)^b$; $n_0 = 1000$, $A_0 = 10^4$ Å$^2$, $b = 1$). Each sampled point defines the centre of a negative microenvironment, encoded identically to positive interface environments. Sampling parameters were calibrated by visual inspection of the point distributions overlaid on representative structures, as shown here.

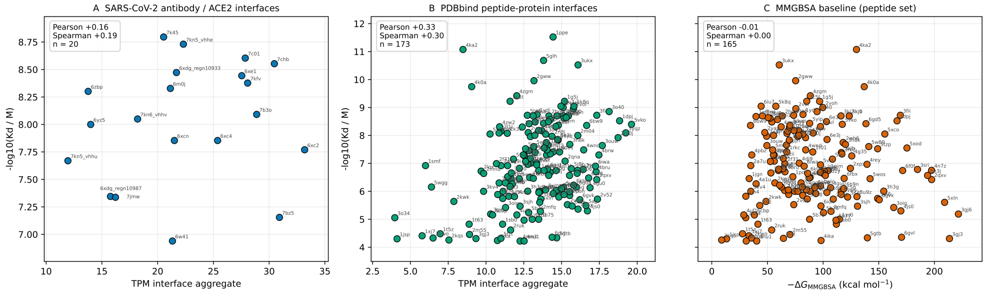


**Fig. A2 The unweighted interface-summed TPM score shows a modest association with absolute binding affinity.** For each interface, anchors with predicted compatibility $\geq 0.5$ were grouped by residue, averaged within each residue, and the per-residue means summed to a single interface score. No per-shard weighting, interface-area normalization or shard exclusion was applied. This score is plotted against experimental binding affinity, expressed as $-\log_{10}(\text{affinity}/1\,\text{M})$ (higher = tighter). **(A)** SARS-CoV-2 antibody/nanobody and ACE2–RBD interfaces ($n = 20$; Pearson $r = 0.16$, Spearman $\rho = 0.19$), for which the correlation is not statistically significant. **(B)** PDBbind peptide–protein interfaces ($n = 173$, Pearson $r = 0.33$, Spearman $\rho = 0.30$), a weak but nonzero positive association. Affinities pool $K_d$, $K_i$ and $\text{IC}_{50}$ measurements reported as exact values. **(C)** A physics-based MM/GBSA baseline on the same peptide-protein complexes ($n = 164$). Pearson $r = 0.33$, Spearman $\rho = 0.30$), computed from 10 ns of explicit-solvent molecular dynamics per complex and plotted as $-\Delta G_{\text{MM/GBSA}}$ so that, as in (A) and (B), larger values correspond to more favorable predicted binding. The MM/GBSA estimate shows no stronger rank association with experimental affinity than the unweighted TPM score, despite being designed specifically for binding free-energy estimation. Absolute MM/GBSA magnitudes (tens to hundreds of $\text{kcal}\,\text{mol}^{-1}$) are dominated by large, incompletely cancelling electrostatic and polar-solvation terms and are not physical binding free energies. Only the rank association with experiment is interpretable here.

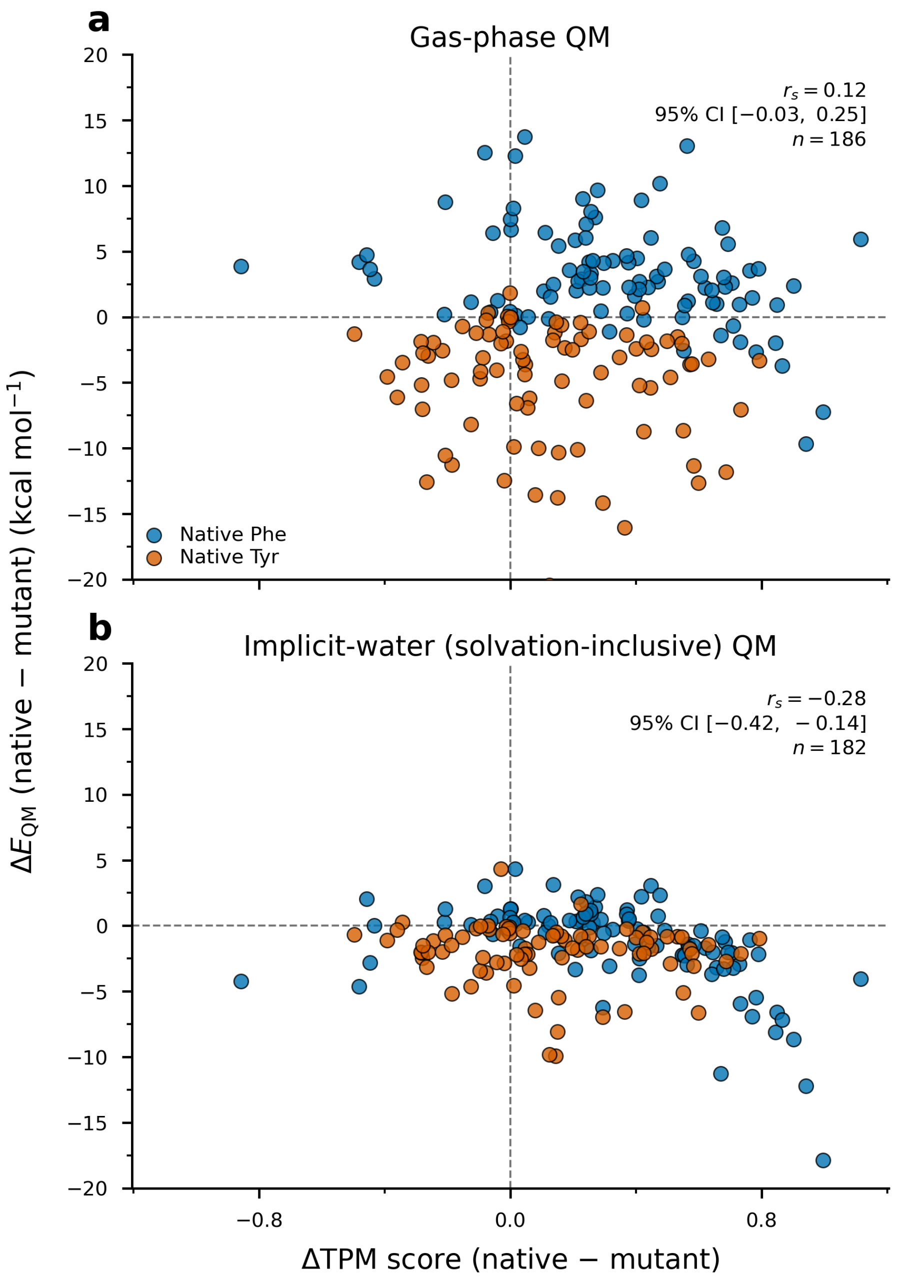


**Fig. A3 The ΔTPM–QM association is specific to solvation-inclusive energetics.** For matched phenylalanine/tyrosine substitutions on the independent peptide-protein set, the difference in summed TPM scores between native and mutant (ΔTPM, native − mutant) is compared with the corresponding quantum-mechanical interaction-energy difference ($\Delta E_{\mathrm{QM}}$, native − mutant; GFN2-xTB). Points are coloured by native residue. **a,** With solvation removed (gas-phase interaction energies), ΔTPM and $\Delta E_{\mathrm{QM}}$ are uncorrelated (Spearman $r_s$ = +0.12, 95% CI [−0.03, 0.25], $n$ = 182). **b,** Adding implicit-water solvation recovers a significant negative monotonic association ($r_s$ = −0.28, 95% CI [−0.42, −0.14], $n$ = 182). The effect is modest in magnitude, as expected for a single-interaction QM proxy compared against a learned interface score, but the sign and significance are specific to the solvation-inclusive calculation since the correlation is absent in gas phase and emerges only once desolvation costs are included. This is consistent with TPMs having implicitly learned solvation-inclusive interface preferences from native, solvated training structures, such that ΔTPM tracks a quantity closer to a net binding contribution than to a raw gas-phase interaction energy. The direct quantitative support for this interpretation is the transfer to solvation- and entropy-inclusive experimental $\Delta\Delta G$ ranking on SKEMPI.

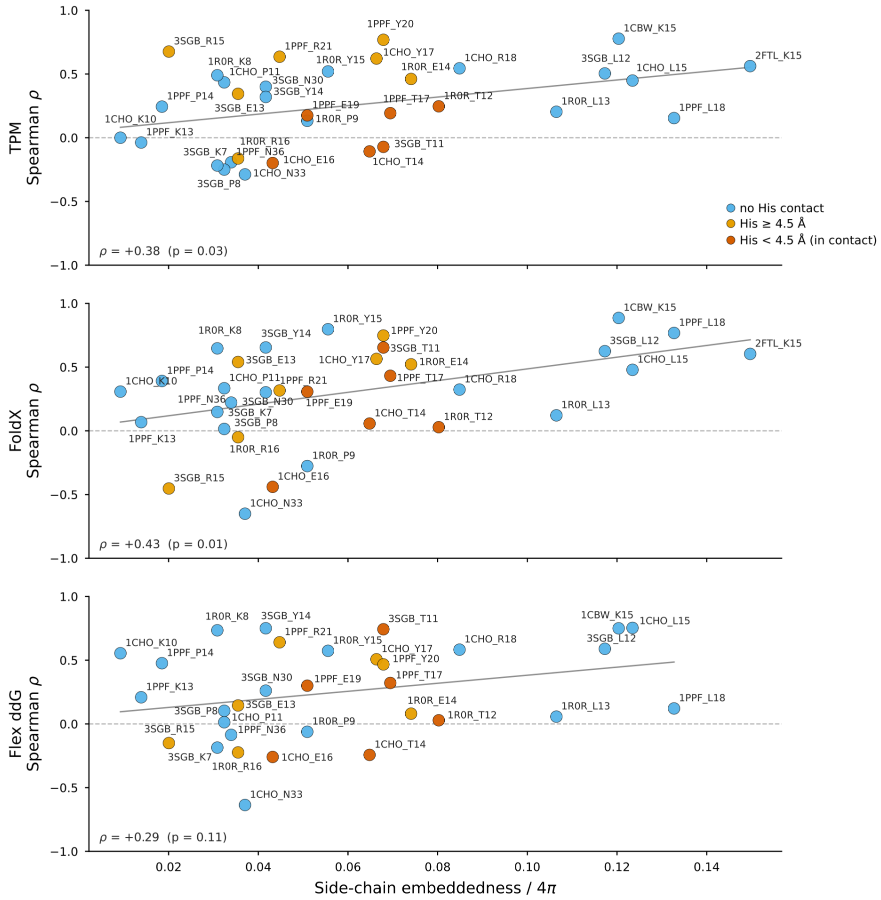


**Fig. A4 Side-chain embeddedness of the ranking pockets.** Per-pocket side-chain embeddedness, defined as the occluded solid-angle fraction of the native side-chain anchor atoms, for the 33 qualifying SKEMPI ranking pockets. Pockets in which a partner histidine lies in direct contact ($< 4.5$ Å) with the scored position are marked, and tend to fall among the lower-ranking pockets for TPM. This is consistent with the present histidine representation not resolving protonation and tautomer state. For zoom-ins on individual pockets and visual illustration of the embeddedness metric see Extended Data Figs. A5–A25.

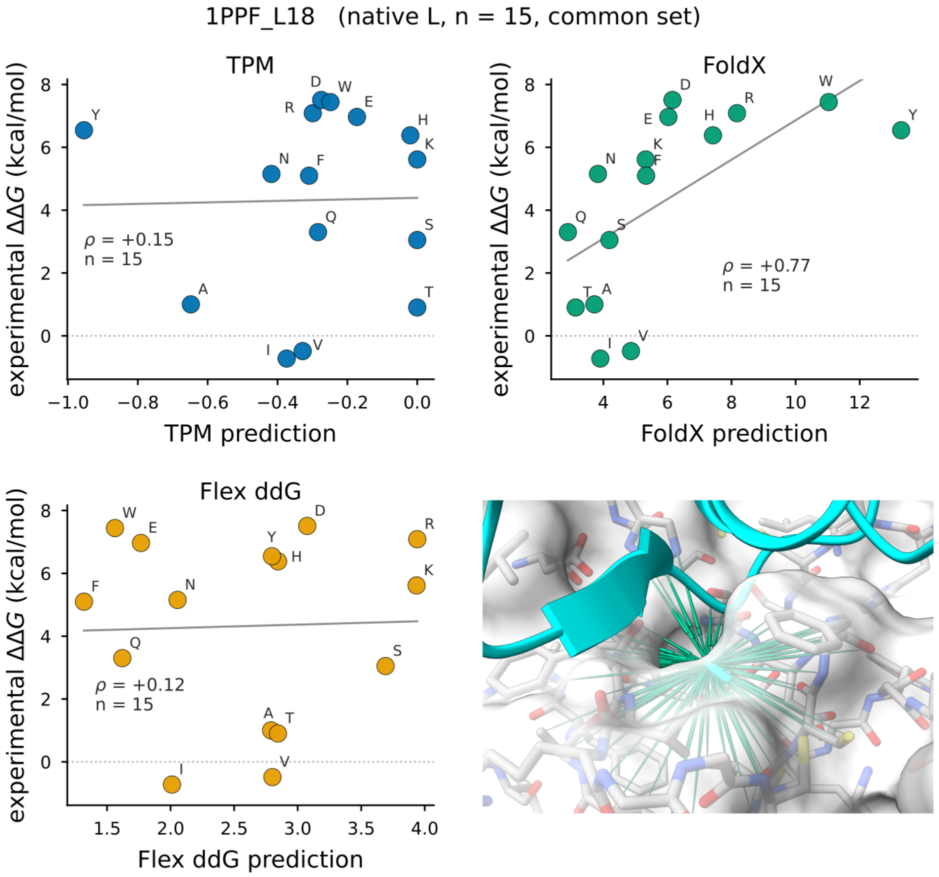


**Fig. A5 Ranking pocket.** Zoom-in of the TPM substitution-preference fields for a single interface pocket from the SKEMPI ranking benchmark (Fig. 4), showing the native side chain and the candidate-substitution fields at this position.

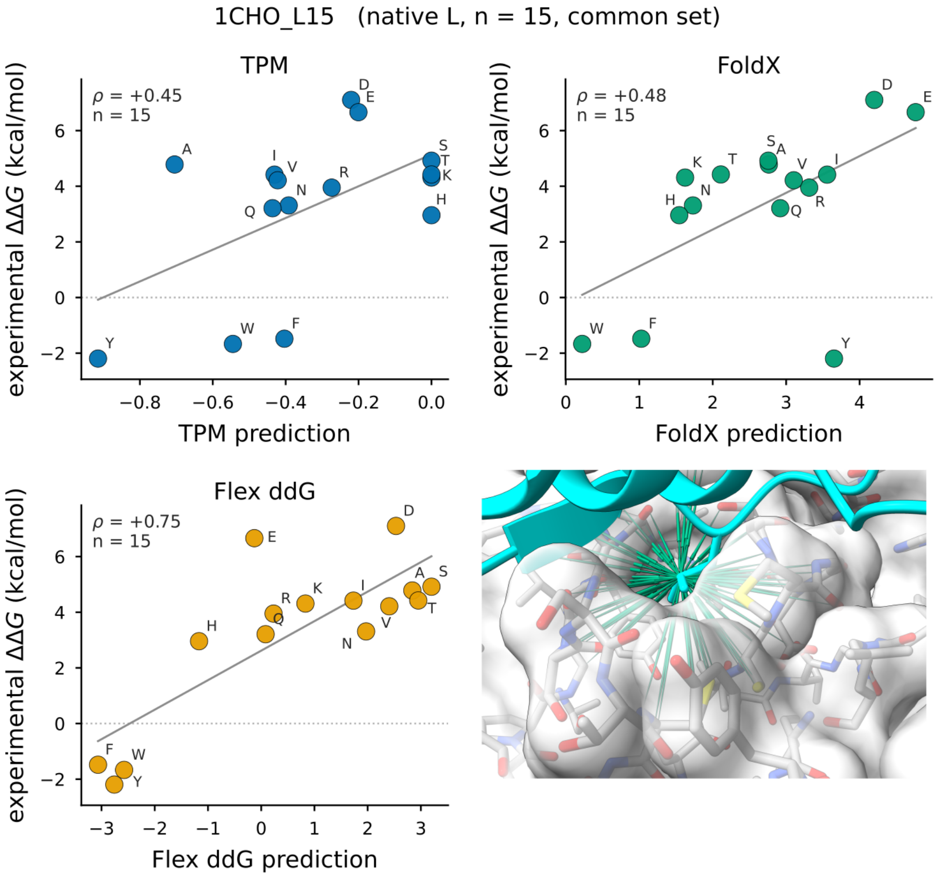


**Fig. A6** **Ranking pocket.** Zoom-in of the TPM substitution-preference fields for a single interface pocket from the SKEMPI ranking benchmark (Fig. 4), showing the native side chain and the candidate-substitution fields at this position.

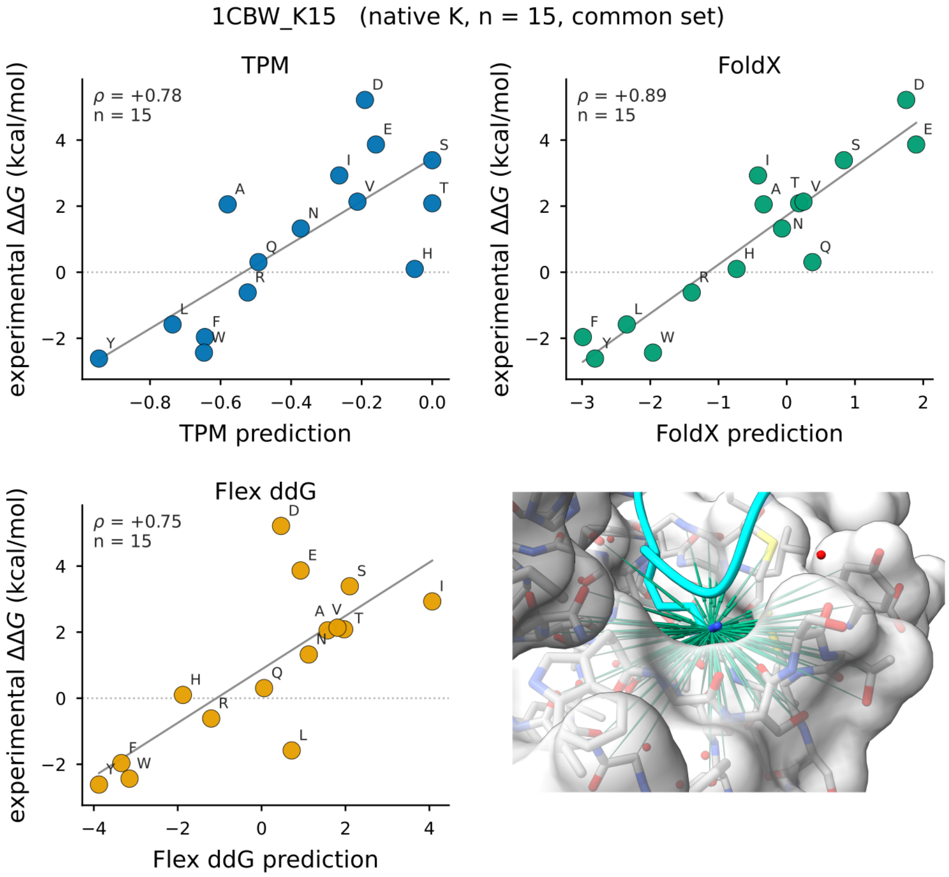


**Fig. A7 Ranking pocket.** Zoom-in of the TPM substitution-preference fields for a single interface pocket from the SKEMPI ranking benchmark (Fig. 4), showing the native side chain and the candidate-substitution fields at this position.

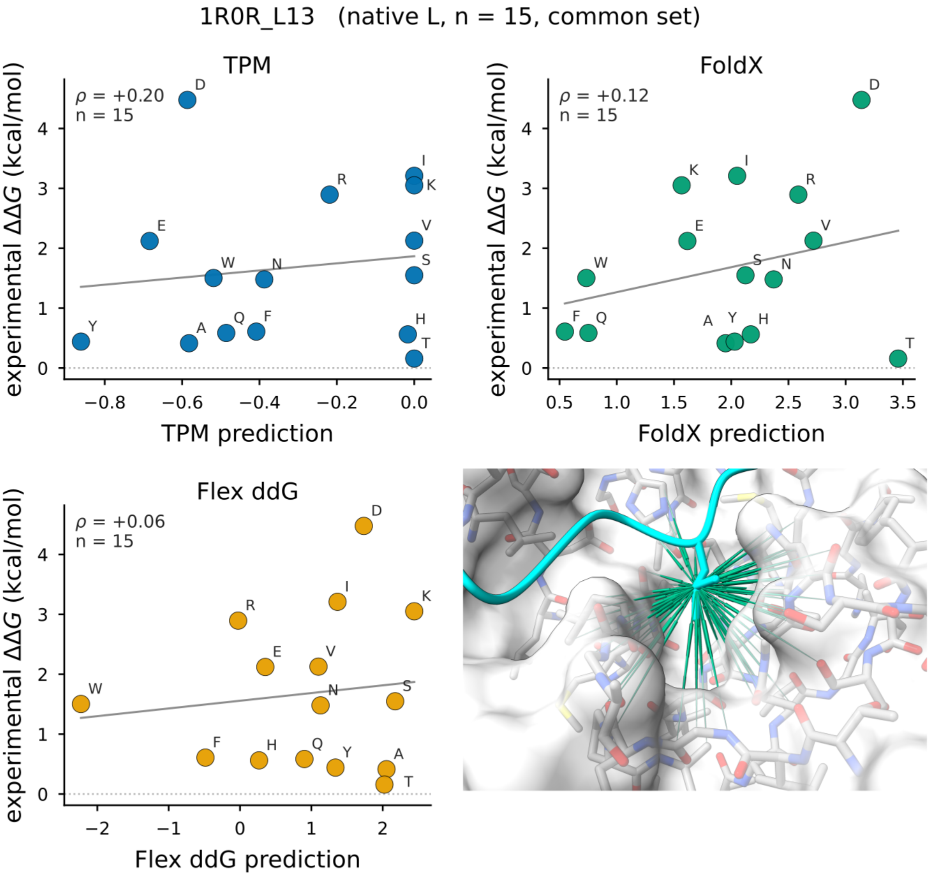


**Fig. A8 Ranking pocket.** Zoom-in of the TPM substitution-preference fields for a single interface pocket from the SKEMPI ranking benchmark (Fig. 4), showing the native side chain and the candidate-substitution fields at this position.

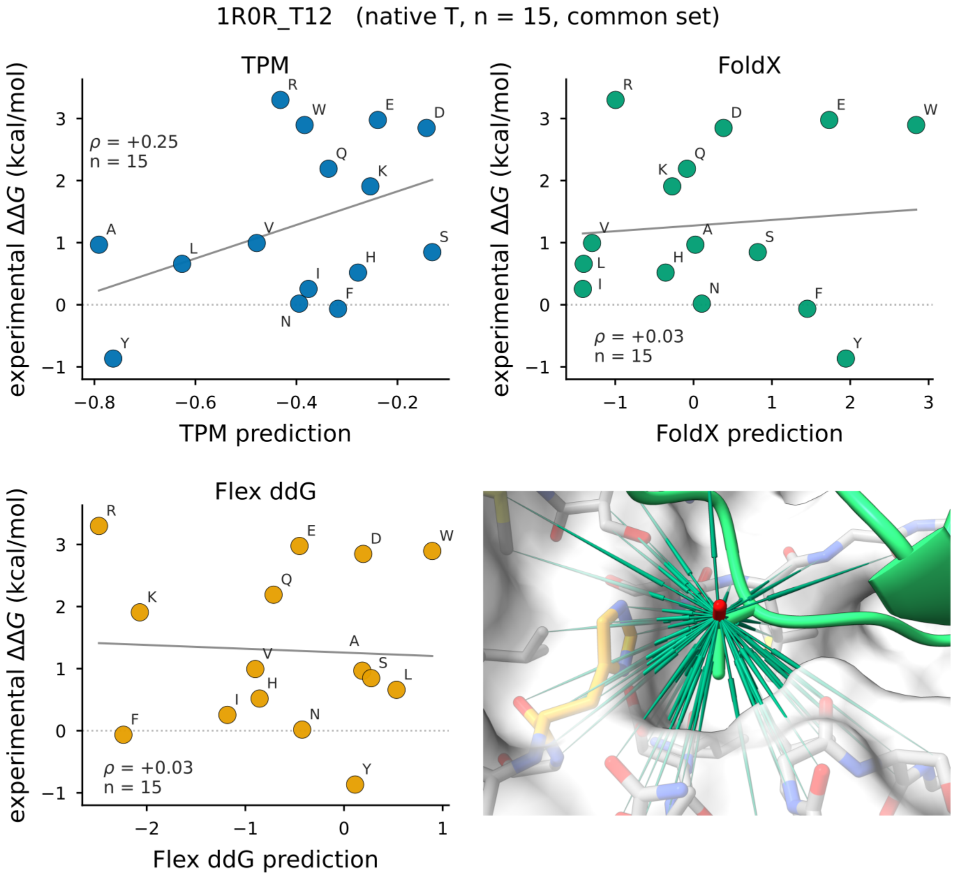


**Fig. A9** **Ranking pocket.** Zoom-in of the TPM substitution-preference fields for a single interface pocket from the SKEMPI ranking benchmark (Fig. 4), showing the native side chain and the candidate-substitution fields at this position.

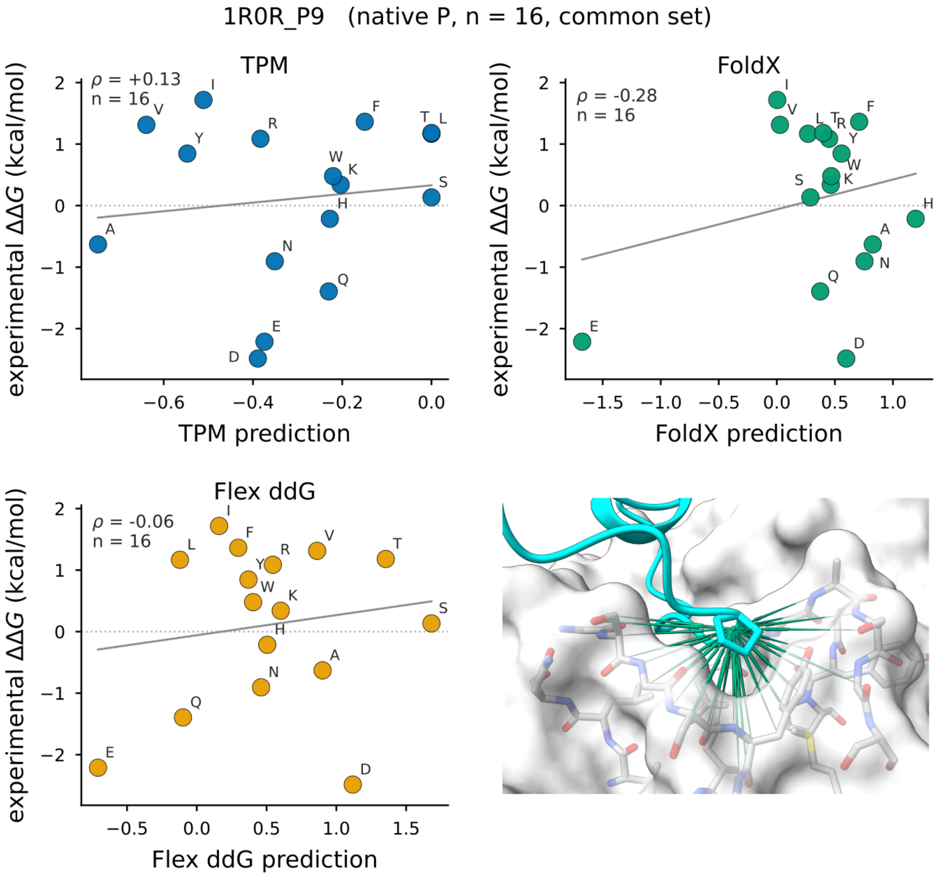


**Fig. A10 Ranking pocket.** Zoom-in of the TPM substitution-preference fields for a single interface pocket from the SKEMPI ranking benchmark (Fig. 4), showing the native side chain and the candidate-substitution fields at this position.

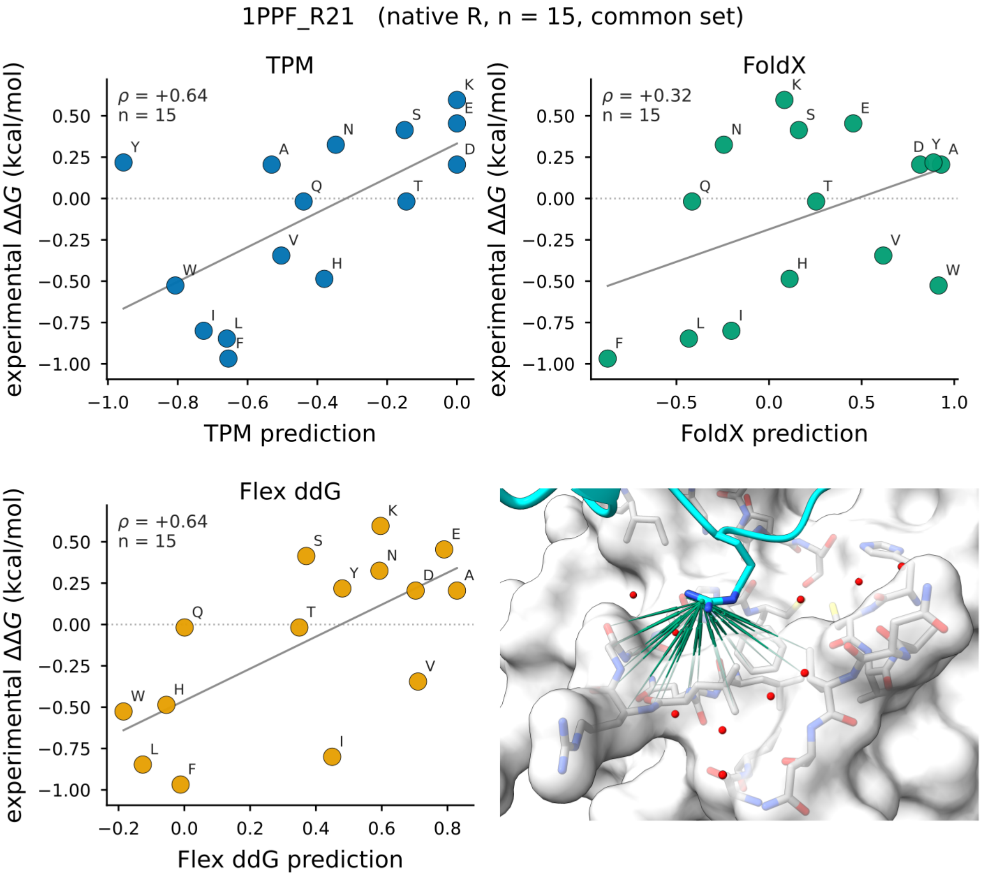


**Fig. A11 Ranking pocket.** Zoom-in of the TPM substitution-preference fields for a single interface pocket from the SKEMPI ranking benchmark (Fig. 4), showing the native side chain and the candidate-substitution fields at this position.

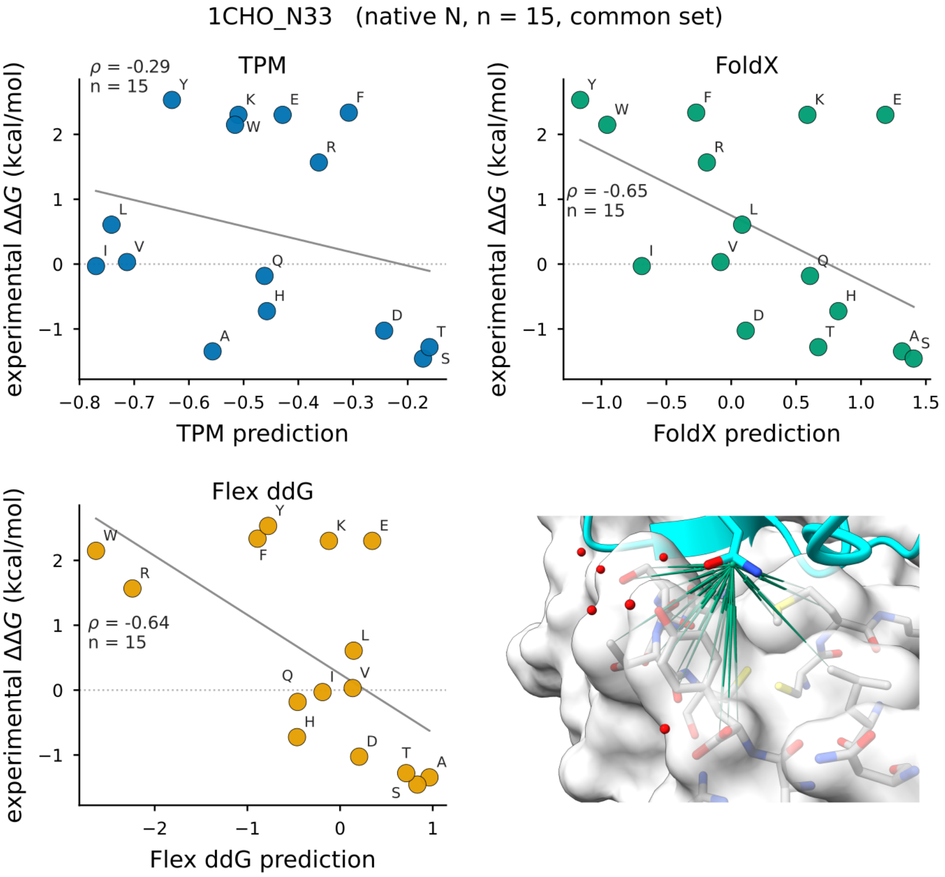


**Fig. A12 Ranking pocket.** Zoom-in of the TPM substitution-preference fields for a single interface pocket from the SKEMPI ranking benchmark (Fig. 4), showing the native side chain and the candidate-substitution fields at this position.

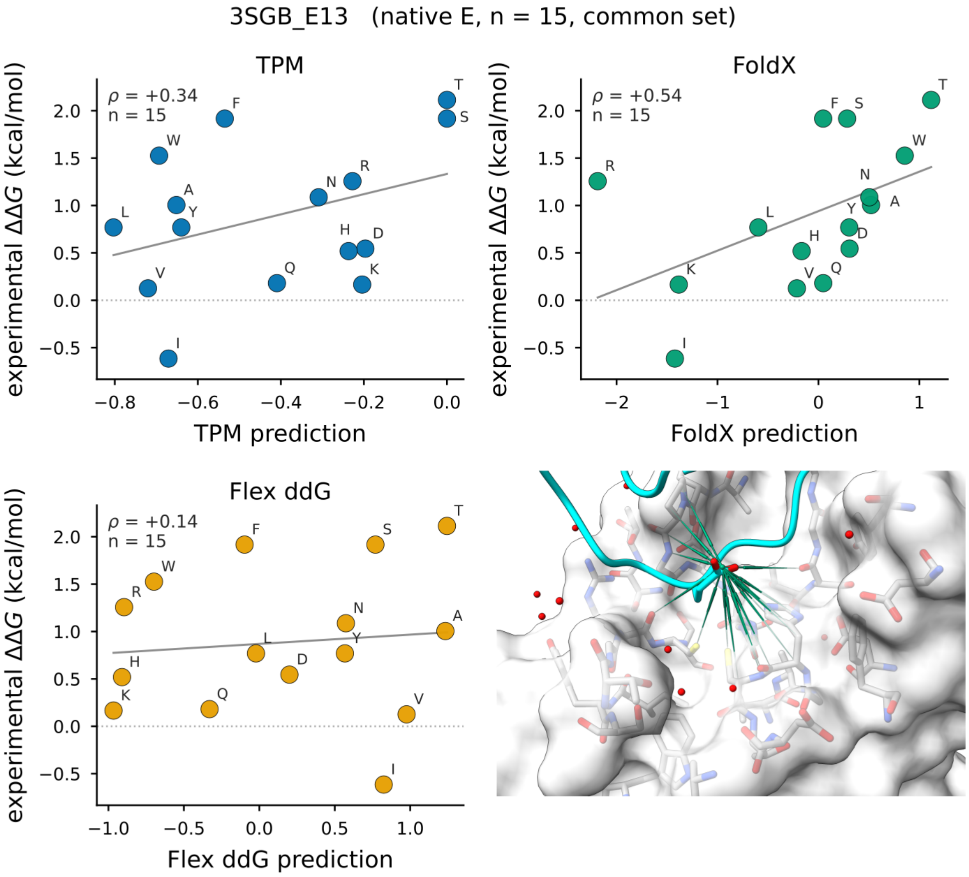


**Fig. A13 Ranking pocket.** Zoom-in of the TPM substitution-preference fields for a single interface pocket from the SKEMPI ranking benchmark (Fig. 4), showing the native side chain and the candidate-substitution fields at this position.

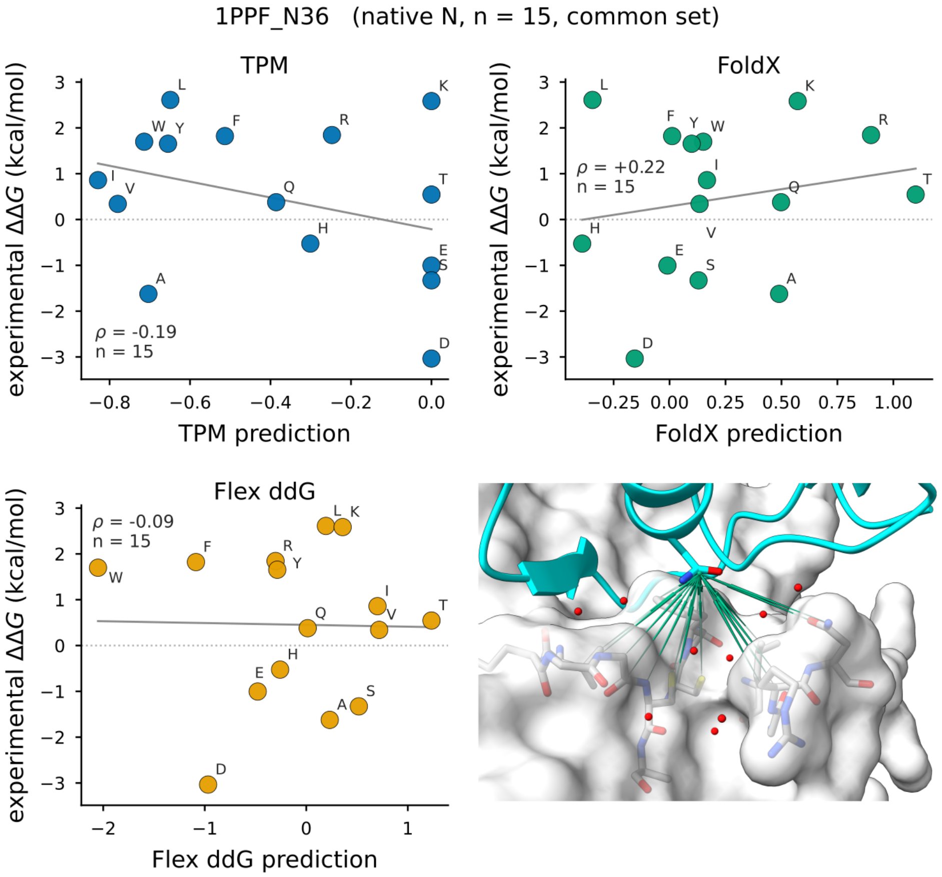


**Fig. A14 Ranking pocket.** Zoom-in of the TPM substitution-preference fields for a single interface pocket from the SKEMPI ranking benchmark (Fig. 4), showing the native side chain and the candidate-substitution fields at this position.

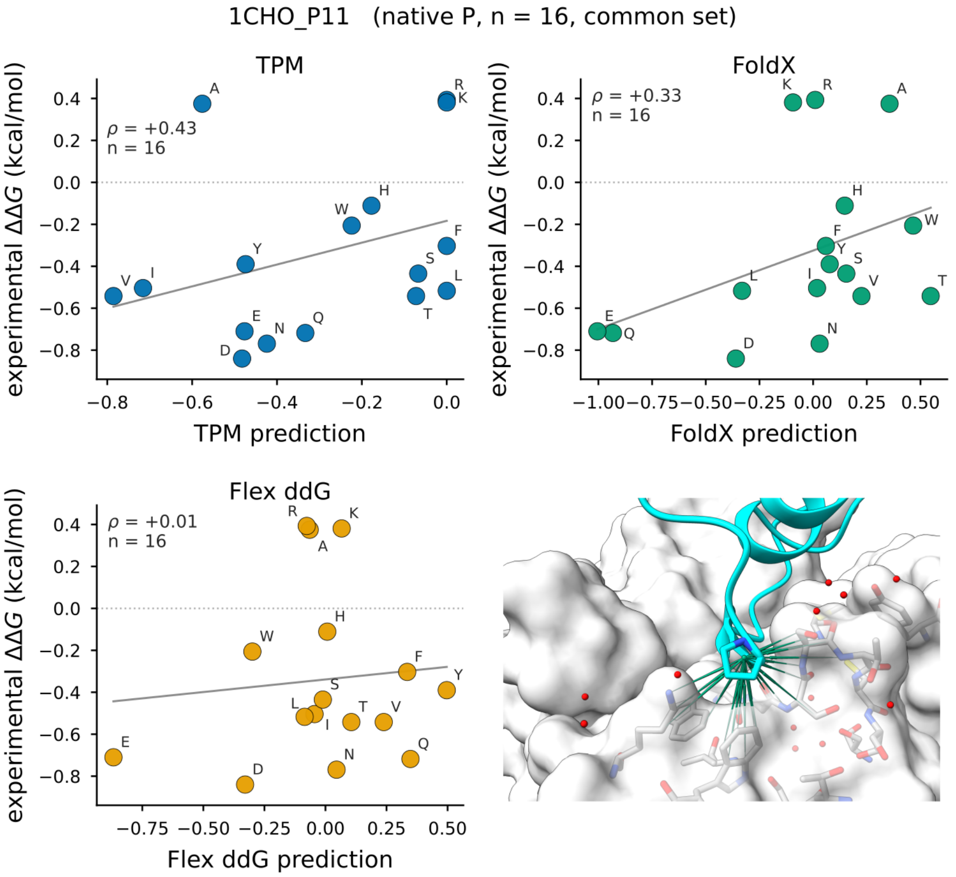


**Fig. A15** **Ranking pocket.** Zoom-in of the TPM substitution-preference fields for a single interface pocket from the SKEMPI ranking benchmark (Fig. 4), showing the native side chain and the candidate-substitution fields at this position.

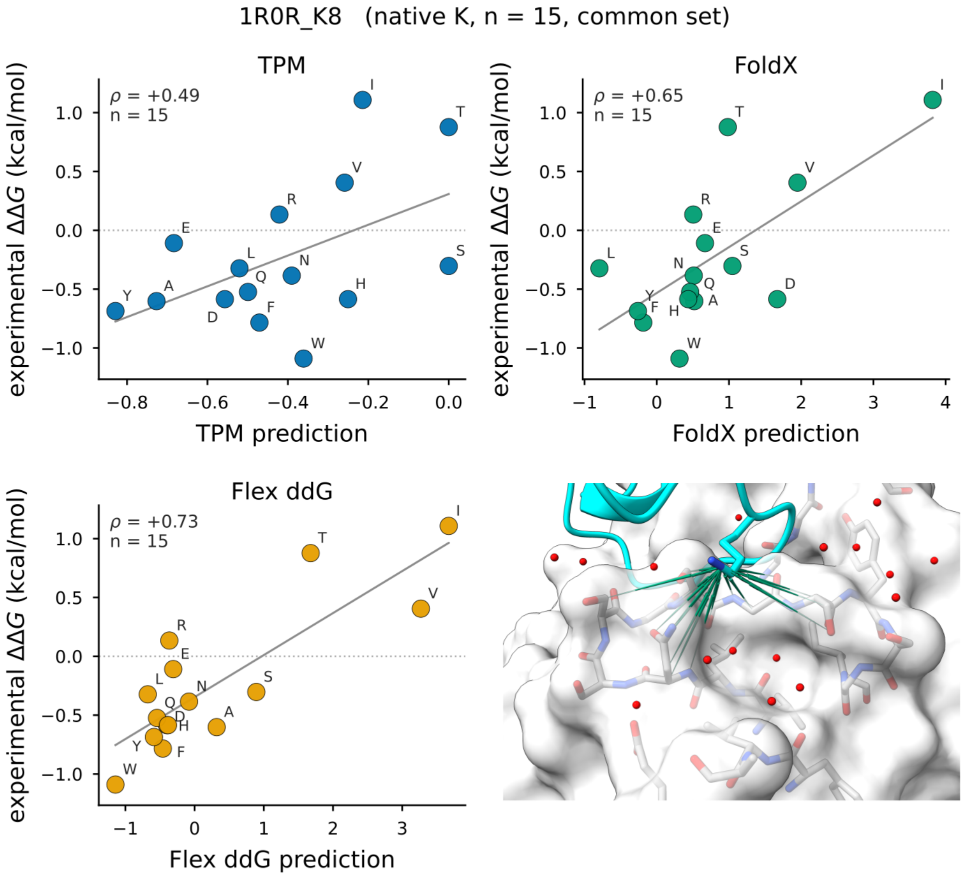


**Fig. A16 Ranking pocket.** Zoom-in of the TPM substitution-preference fields for a single interface pocket from the SKEMPI ranking benchmark (Fig. 4), showing the native side chain and the candidate-substitution fields at this position.

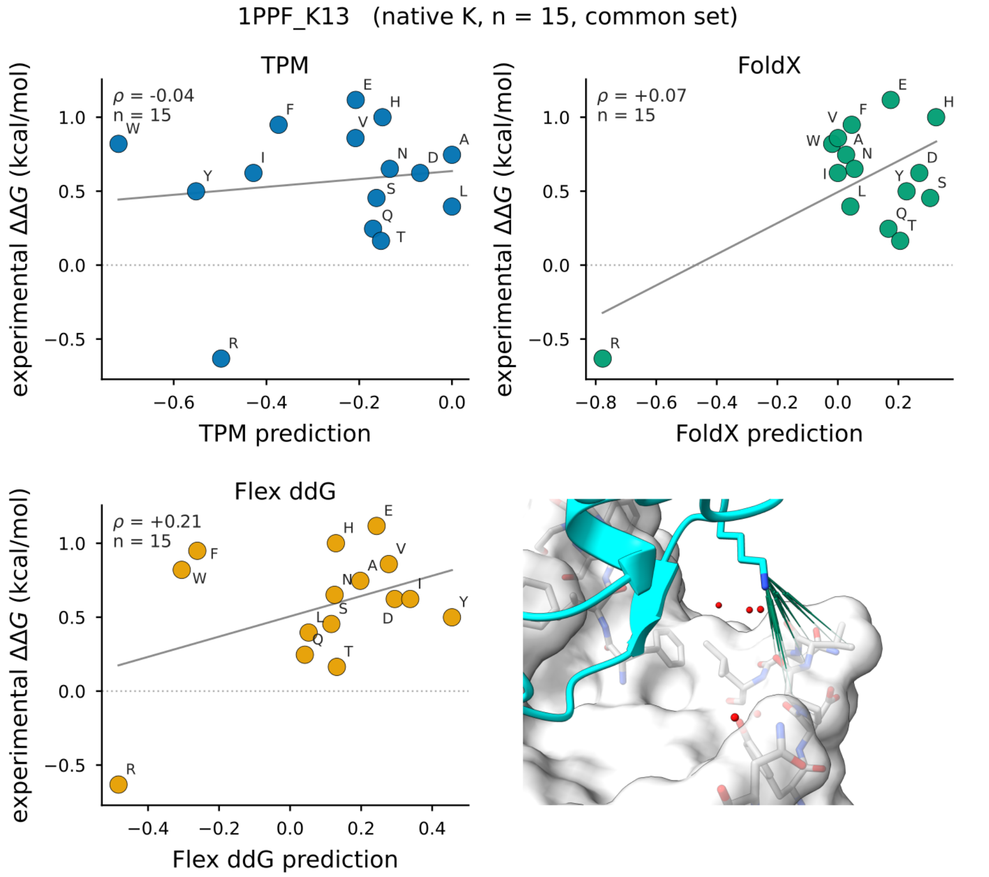


**Fig. A17** **Ranking pocket.** Zoom-in of the TPM substitution-preference fields for a single interface pocket from the SKEMPI ranking benchmark (Fig. 4), showing the native side chain and the candidate-substitution fields at this position.

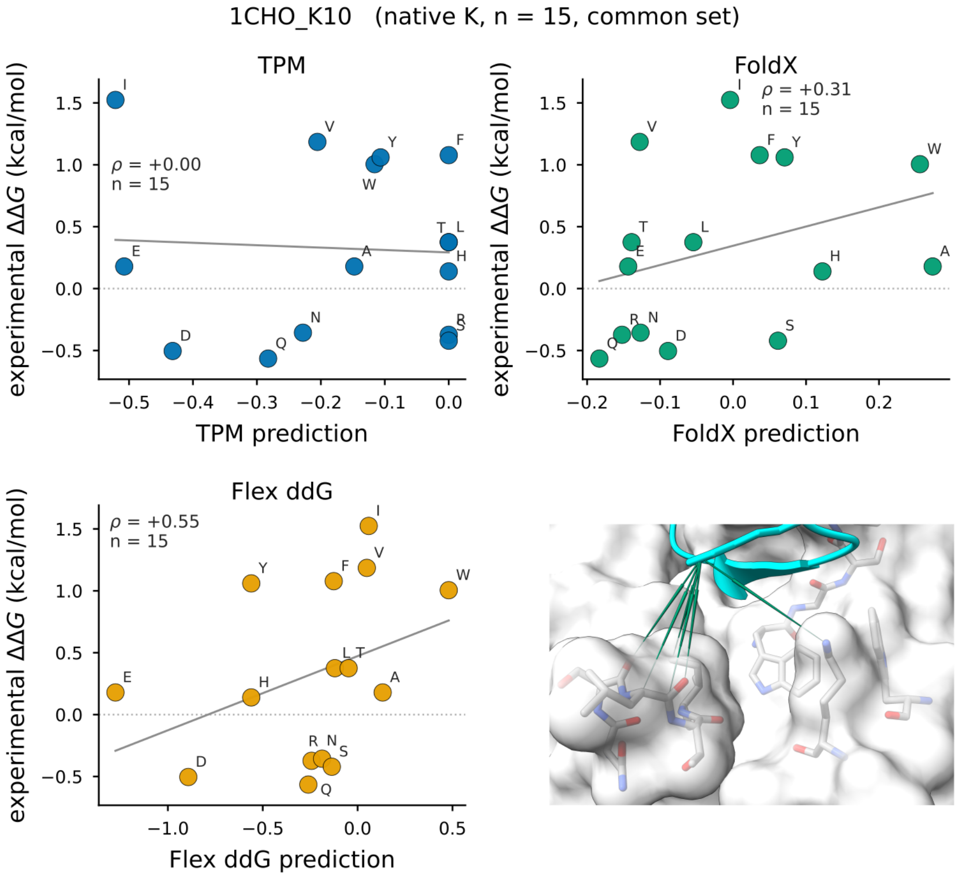


**Fig. A18** **Ranking pocket.** Zoom-in of the TPM substitution-preference fields for a single interface pocket from the SKEMPI ranking benchmark (Fig. 4), showing the native side chain and the candidate-substitution fields at this position.

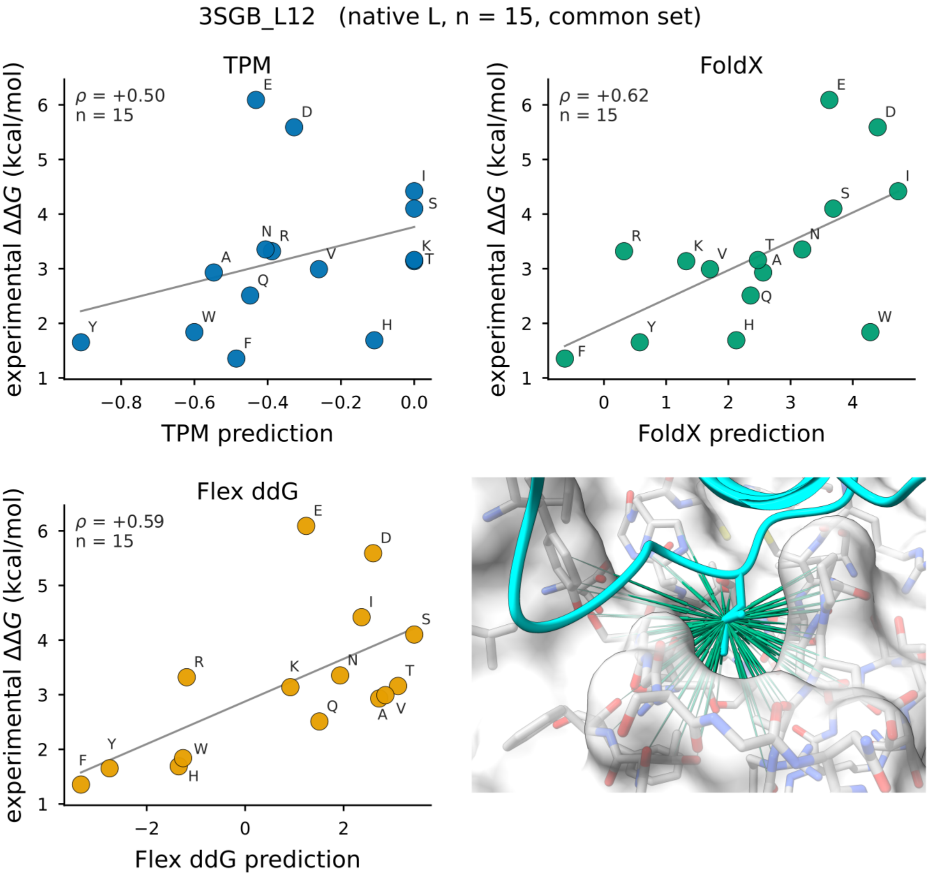


**Fig. A19 Ranking pocket.** Zoom-in of the TPM substitution-preference fields for a single interface pocket from the SKEMPI ranking benchmark (Fig. 4), showing the native side chain and the candidate-substitution fields at this position.

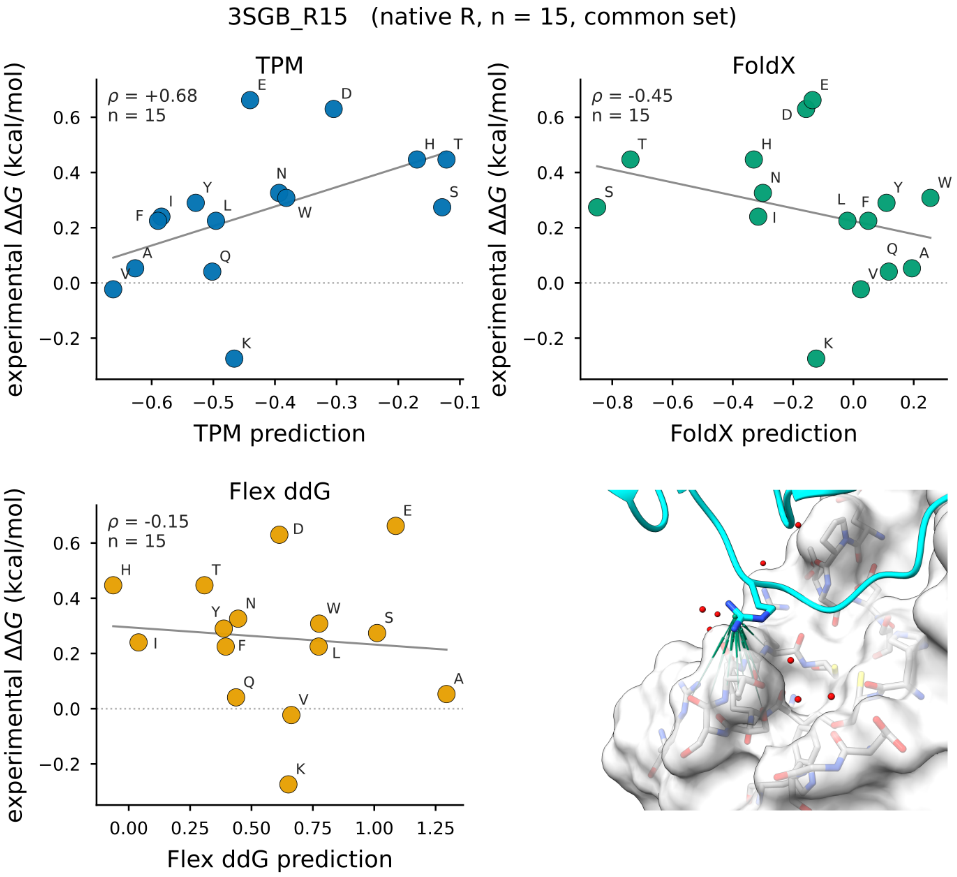


**Fig. A20 Ranking pocket.** Zoom-in of the TPM substitution-preference fields for a single interface pocket from the SKEMPI ranking benchmark (Fig. 4), showing the native side chain and the candidate-substitution fields at this position.

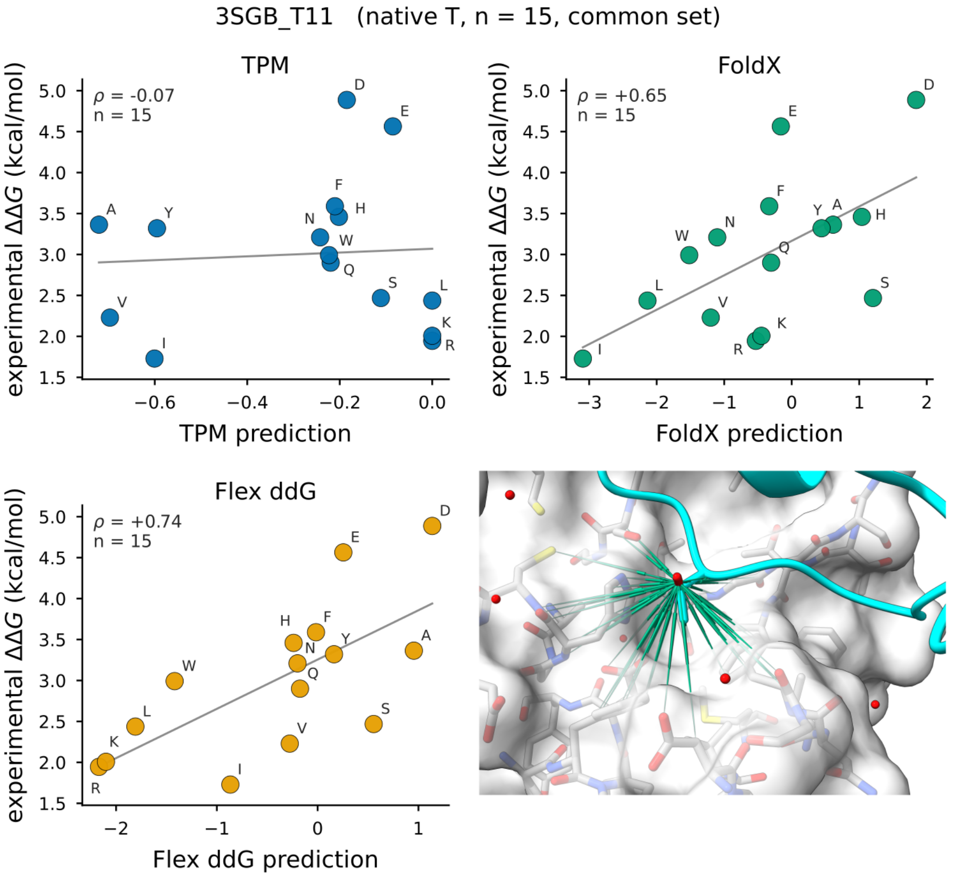


**Fig. A21** **Ranking pocket.** Zoom-in of the TPM substitution-preference fields for a single interface pocket from the SKEMPI ranking benchmark (Fig. 4), showing the native side chain and the candidate-substitution fields at this position.

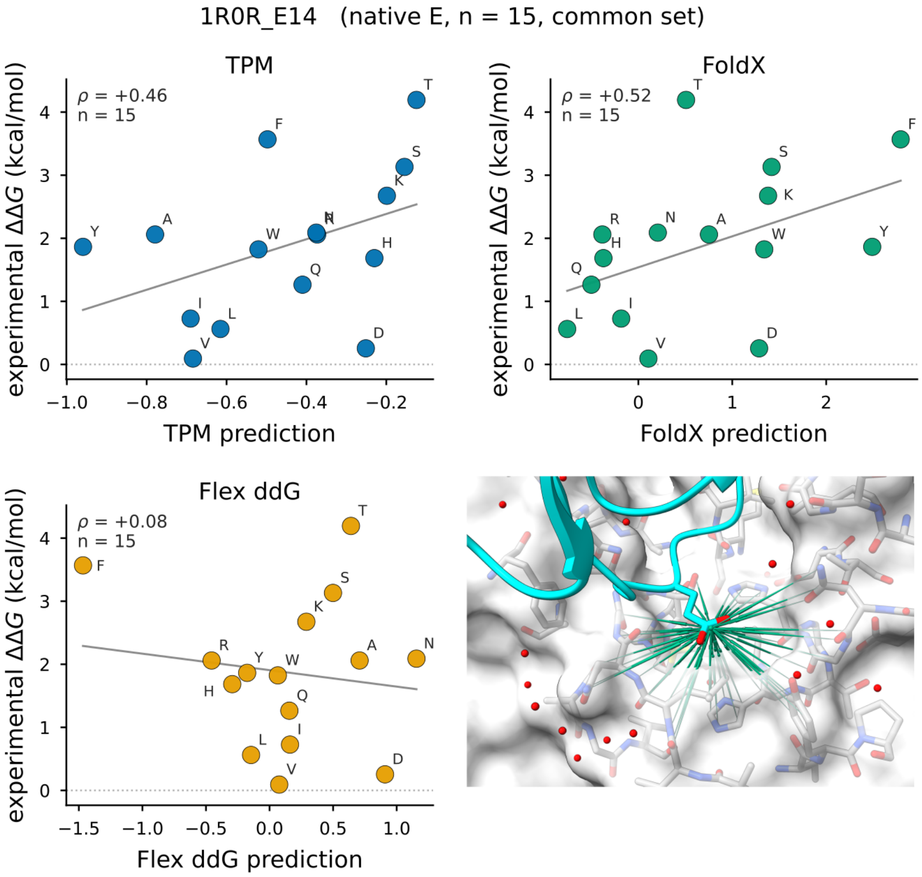


**Fig. A22 Ranking pocket.** Zoom-in of the TPM substitution-preference fields for a single interface pocket from the SKEMPI ranking benchmark (Fig. 4), showing the native side chain and the candidate-substitution fields at this position.

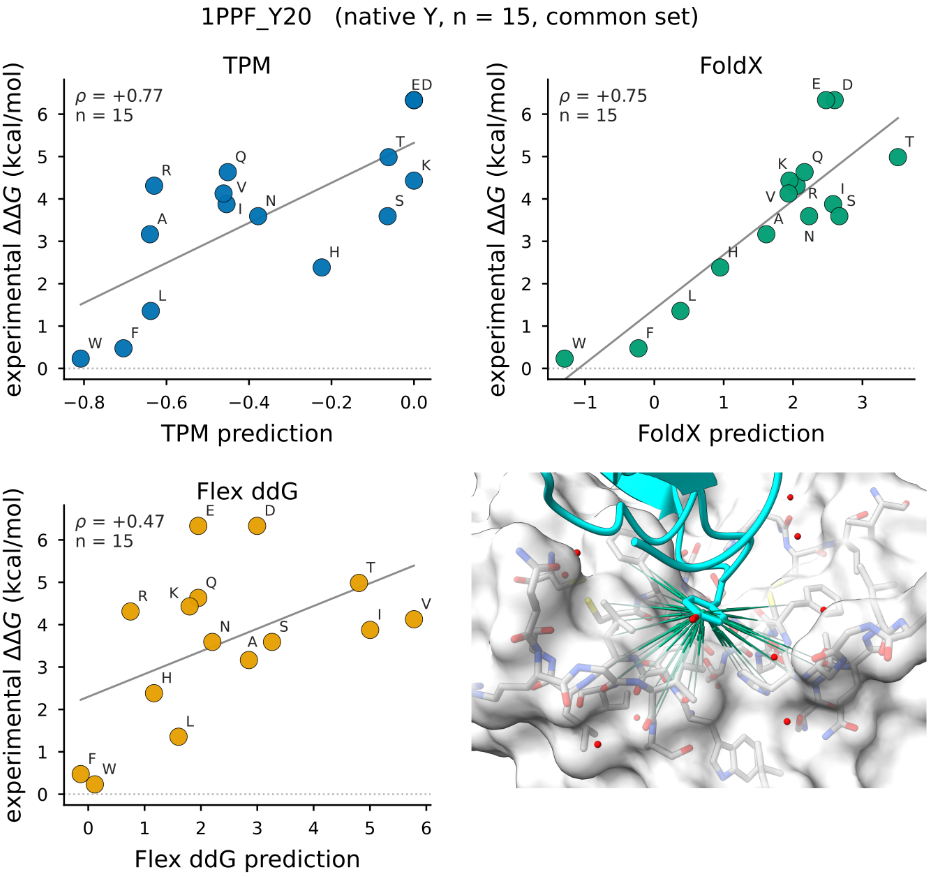


**Fig. A23 Ranking pocket.** Zoom-in of the TPM substitution-preference fields for a single interface pocket from the SKEMPI ranking benchmark (Fig. 4), showing the native side chain and the candidate-substitution fields at this position.

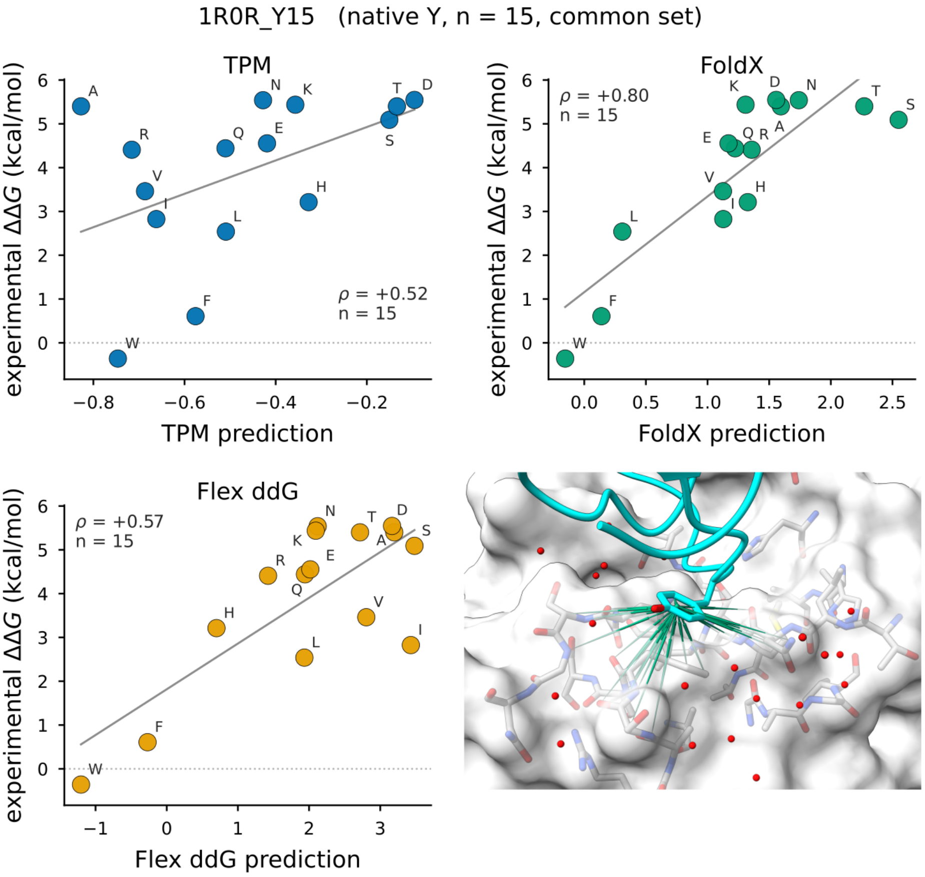


**Fig. A24 Ranking pocket.** Zoom-in of the TPM substitution-preference fields for a single interface pocket from the SKEMPI ranking benchmark (Fig. 4), showing the native side chain and the candidate-substitution fields at this position.

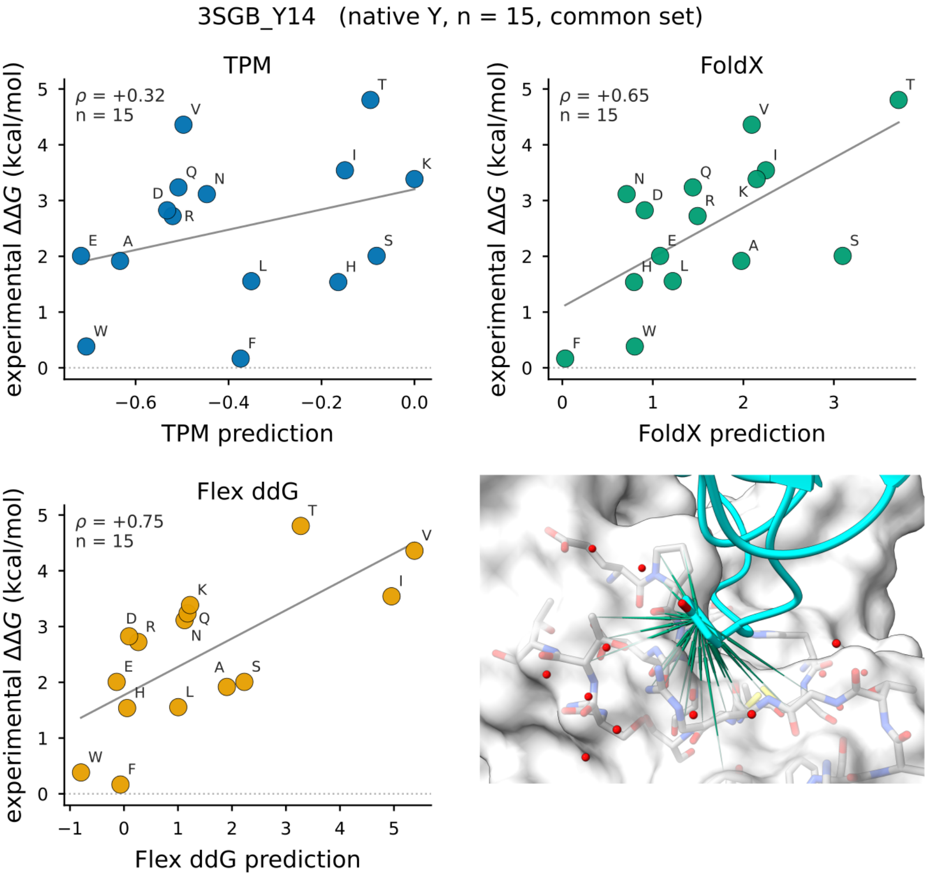


**Fig. A25 Ranking pocket.** Zoom-in of the TPM substitution-preference fields for a single interface pocket from the SKEMPI ranking benchmark (Fig. 4), showing the native side chain and the candidate-substitution fields at this position.
