## Supplementary figures and images for "Local structural preference maps encode transferable protein-interface energetics"

### Extended Data Video 1

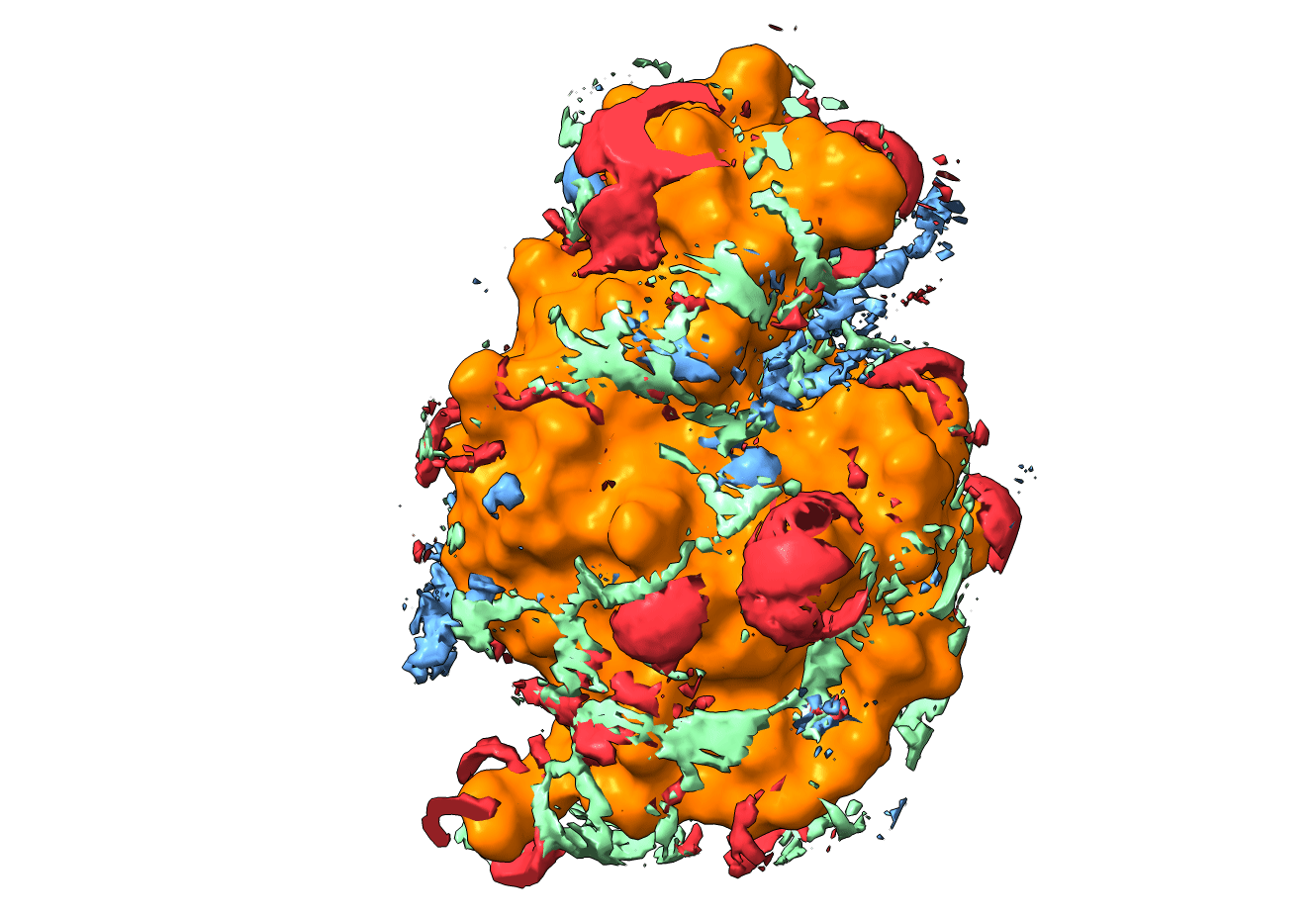
